# Activity-dependent development of the body’s touch receptors

**DOI:** 10.1101/2023.09.23.559109

**Authors:** Celine Santiago, Nikhil Sharma, Nusrat Africawala, Julianna Siegrist, Annie Handler, Aniqa Tasnim, Rabia Anjum, Josef Turecek, Brendan P. Lehnert, Sophia Renauld, Michael Nolan-Tamariz, Michael Iskols, Alexandra R. Magee, Suzanne Paradis, David D. Ginty

## Abstract

We report a role for activity in the development of the primary sensory neurons that detect touch. Genetic deletion of Piezo2, the principal mechanosensitive ion channel in somatosensory neurons, caused profound changes in the formation of mechanosensory end organ structures and altered somatosensory neuron central targeting. Single cell RNA sequencing of *Piezo2* conditional mutants revealed changes in gene expression in the sensory neurons activated by light mechanical forces, whereas other neuronal classes were less affected. To further test the role of activity in mechanosensory end organ development, we genetically deleted the voltage-gated sodium channel Na_v_1.6 (*Scn8a*) in somatosensory neurons throughout development and found that *Scn8a* mutants also have disrupted somatosensory neuron morphologies and altered electrophysiological responses to mechanical stimuli. Together, these findings indicate that mechanically evoked neuronal activity acts early in life to shape the maturation of the mechanosensory end organs that underlie our sense of gentle touch.

## Introduction

The sense of touch is one of the earliest to develop, and alterations in tactile sensory inputs during early postnatal life cause short-term effects on physiology and long-term changes in animal behavior ^1-4^. However, the molecular, cellular, and circuit-level mechanisms by which touch-evoked activity influences the maturation of the nervous system remain poorly understood. In particular, while classic limb amputation, whisker plucking, and nerve lesion experiments demonstrated that peripheral inputs shape the development of somatosensory cortex and other central brain regions ^5-8^, it remains an open question whether and how neuronal activity influences the maturation of the primary somatosensory neurons that detect tactile stimulation.

Light touch sensation is mediated by low threshold mechanoreceptors (LTMRs), sensory neurons whose cell bodies reside in the dorsal root ganglia (DRG) or trigeminal ganglia and that form bifurcated axons, with one branch traveling to the periphery, where the axon is activated by mechanical stimuli, and the other forming synapses in the central nervous system. The peripheral axons of LTMRs associate with specialized non-neuronal cells to form complex multicellular structures known as mechanosensory end organs. These end organs, which are found across the skin and in deeper tissues throughout the body, have distinct functional response properties which arise in part from their specialized morphologies ^9^. Meissner corpuscles, which are found in the glabrous (non-hairy) skin of mammals, are sensitive to skin indentation and low frequency vibration (∼10-100 Hz), and are composed of 1-2 mechanosensory axons that form close associations with specialized glial cells known as lamellar cells ^10^. Pacinian corpuscles, large mechanically sensitive structures that consist of a single axon wrapped by multiple layers of non-neuronal cells, are tuned to high frequency vibration (∼40-1000 Hz) and in rodents are enriched in the interosseous membranes surrounding bones, though in primates they are also present in glabrous skin ^11^. In hairy skin, multiple genetically, functionally, and anatomically distinct classes of LTMRs form specialized axonal endings around hair follicles ^12-15^, rendering them exquisitely sensitive to skin indentation and hair deflection. The mechanisms that underlie the development of LTMRs and the peripheral mechanosensory end organs they innervate have been studied using detailed anatomical and molecular analyses ^16,17^. However, how neuronal activity influences the establishment of touch sensory end organs and their specialized morphologies remains unknown.

In mammals, Piezo2 (piezo-type mechanosensitive ion channel component 2) is required to transduce mechanical stimuli into electrical activity in mature somatosensory neurons and plays an essential role in both exteroception and interoception. In addition to its function in the detection of innocuous touch and some forms of painful touch ^18-23^, Piezo2 is critical for normal proprioception ^24^, breathing ^25^, blood pressure regulation ^26,27^, urination ^28^, sexual function ^29^, gut motility ^30^, and colon sensation ^31^. The finding that loss of Piezo2 results in the near abolishment of responses to touch in the DRG or trigeminal ganglia ^18,23,32,33^ offers an entry point for exploring the role of activity during somatosensory system maturation. While Piezo2 and the related protein Piezo1 regulate the development of several non-neuronal tissues, including the cardiovascular system (Piezo1) ^34-36^ and the musculoskeletal system (Piezo2) ^37,38^, they have not been reported to influence the development of somatosensory neurons or their downstream circuits.

We found that Piezo2 is required for the structural and transcriptional maturation of LTMRs, the primary sensory neurons that detect gentle touch. *Cdx2-Cre* mediated deletion of Piezo2 in somatosensory neurons throughout development causes disruptions in the structures of peripheral mechanosensory end organs and in the central projection patterns of somatosensory neurons. Single cell RNA sequencing of DRG neurons revealed that while the major transcriptionally defined cell types were present in *Piezo2* conditional mutants, significant changes in gene expression were detected in mechanosensory neuron subtypes. Finally, disruptions in mechanically evoked neuronal activity and peripheral mechanosensory end organs following genetic deletion of the voltage-gated sodium channel Na_v_1.6 (*Scn8a*) suggest a generalizable role for neuronal activity in mechanosensory end organ morphogenesis. Thus, during development, Piezo2 and Na_v_-dependent electrical activity in mechanosensory neurons regulates their complex peripheral morphologies within the end organ structure. Accordingly, tactile experiences during embryonic and postnatal development may influence the formation of the first node in the somatosensory pathway by controlling maturation of peripheral touch receptors.

## Results

### Piezo2 is required for normal development of peripheral mechanosensory end organs

We previously showed using electrophysiological recordings that responses to low force mechanical stimuli are not detected in lumbar-level dorsal root ganglia (DRG) of *Cdx2-Cre/+; Piezo2^flox/Null^ or Cdx2-Cre/+; Piezo2^flox/flox^* animals (hereafter referred to as *Cdx2-Cre Piezo2 cKO*)^32,33^. Responses to optical stimulation of the paw in mutants expressing the light-activated cation channel ReaChR are still present, indicating that sensory neurons in these mutants innervate the skin and can fire action potentials when their peripheral terminals are activated, in this case by light ^32^. To examine peripheral innervation in *Cdx2-Cre Piezo2 cKO* mutants more closely, we collected hind paw glabrous skin, hind limb interosseous membrane, and trunk hairy skin and performed immunohistochemistry to visualize several major mechanoreceptor end organ classes.

We were able to identify the expected mechanosensory peripheral endings in *Cdx2-Cre Piezo2 cKO* animals but detected notable defects in the morphologies of specific end organs. Meissner corpuscles were present in the plantar pads of hind paw glabrous skin in the mutants, but often displayed enlarged and disorganized morphologies, and supernumerary innervation by Neurofilament heavy chain (NFH)^+^ axons (Figure 1A). The enlarged Meissner corpuscles seen in *Cdx2-Cre Piezo2 cKO* mutants also contained more S100^+^ lamellar cells (Supplemental Figure 1A). Despite the changes in morphology observed by light microscopy, high resolution imaging using electron microscopy revealed no defects in the ultrastructural organization of the corpuscles, as Meissner corpuscles of both *Cdx2-Cre Piezo2 cKO* and control animals contained multiple axon profiles tightly wrapped by lamellar cells (Supplemental Figure 1B).

**Figure 1:**
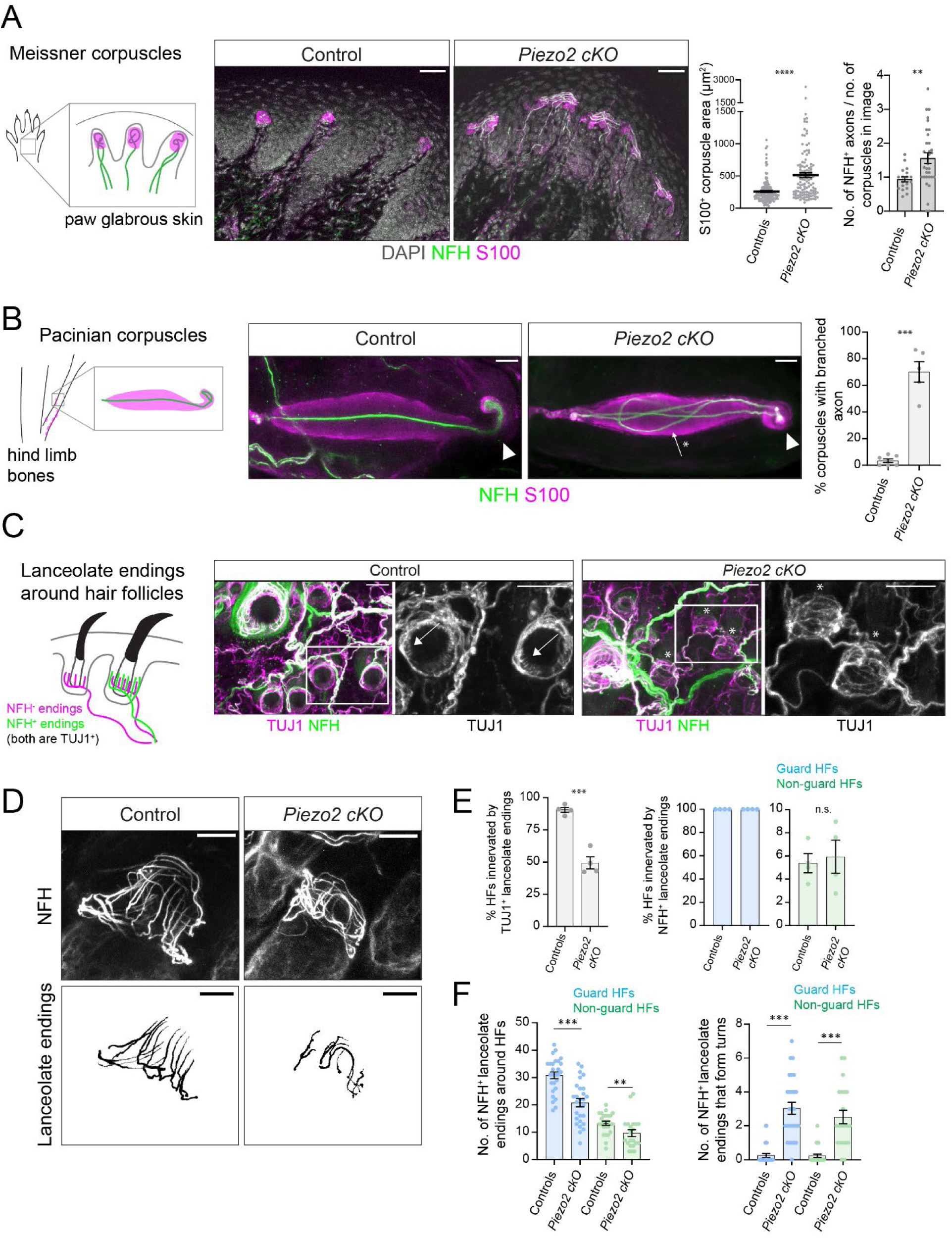
Piezo2 is required for normal mechanosensory end organ structures. **A.** Meissner corpuscles are detected in the plantar pads of hind paw glabrous skin in mice and are labeled by anti-NFH (labels myelinated axons, green) and anti-S100 (labels glial cells, magenta) antibodies. In *Cdx2-Cre; Piezo2 cKO* animals, the S100^+^ areas of Meissner corpuscles are larger (****p<0.0001, Mann-Whitney U test) and the corpuscles are innervated by more NFH^+^ axons than in controls (**p<0.01, Welch’s t test). For area quantification, each data point indicates one Meissner corpuscle; averages are shown by horizontal bars; error bars indicate the standard error of the mean (s.e.m.). For NFH^+^ axon quantification, each data point indicates one image; averages are shown by horizontal bars; error bars indicate the standard error of the mean (s.e.m.). 5 control littermates and 7 *Cdx2-Cre; Piezo2 cKO* animals were analyzed; see Supplemental Table 1 for exact genotypes. Scale bar = 30 µm. See also Figure S1. **B.** Pacinian corpuscles are found in the interosseous membrane surrounding the fibula of the mouse hind limb and are stained using anti-NFH (green) and anti-S100 (magenta) antibodies. In adult controls, almost all Pacinian corpuscles contain a single, linear NFH^+^ axon that selectively branches in the ultraterminal region (indicated by the arrowheads). In *Cdx2-Cre Piezo2 cKO* animals, the majority of Pacinian corpuscles contain an NFH^+^ axon that branches once or multiple times prior to the ultraterminal region (arrow with asterisk). Right: Quantification of the percentage of Pacinian corpuscles with a branched axon (***p<0.001, Welch’s t test). Each data point indicates one animal; averages are plotted; error bars indicate the s.e.m. Scale bar = 20 µm. **C.** Left: Mouse trunk hairy skin is innervated by neurons that form NFH^-^ (magenta) or NFH^+^ (green) lanceolate endings around the base of hair follicles (HFs). Anti-beta III tubulin (TUJ1) staining labels all neurons. Right: Example images of hairy skin. Insets show HFs innervated by TUJ1^+^ lanceolate endings (arrows) or HFs not innervated by TUJ1^+^ lanceolate endings (asterisks). Scale bar = 30 µm. **D.** Example images of NFH^+^ axonal endings around non-guard hair follicles. The lanceolate endings were reconstructed using z stacks; maximum projections of reconstructions are shown below. Scale bar = 20 µm. **E.** Quantification of hair follicle innervation by TUJ1^+^ lanceolate endings (labels all neurons) or NFH^+^ lanceolate endings (labels Aβ-LTMRs). *Cdx2-Cre; Piezo2 cKO* animals display a reduction in the percentage of all hair follicles (HFs) innervated by TUJ1^+^ lanceolate endings (gray bars)(***p<0.001, unpaired t test), but no change in the percentage of guard HFs (blue) or non-guard HFs (green) innervated by NFH^+^ lanceolate endings (unpaired t test). Each data point indicates one animal; averages are plotted; error bars indicate the s.e.m. **F.** Quantification of NFH^+^ axonal structures formed by lanceolate endings around guard HFs (blue) or non-guard HFs (green). Left: For both types of hairs, fewer NFH^+^ lanceolate endings were present in *Cdx2-Cre; Piezo2 cKO* animals (***p<0.0001, unpaired t test for guard HFs, **p<0.001, Mann-Whitney U test for non-guard HFs). Right: NFH^+^ lanceolate endings formed more aberrant turns along the circumferential axis in *Piezo2* mutants (***p<0.0001, Mann-Whitney U test). Each data point indicates one hair follicle; averages are plotted; error bars indicate the s.e.m. Data were collected from 5 control animals and 5 *Cdx2-Cre; Piezo2 cKO* mutants.

Pacinian corpuscles were also present in the expected location and numbers around the fibula of *Cdx2-Cre Piezo2 cKO* animals. However, in mutants they were frequently innervated by an axon that formed a complex, irregular branching pattern, in contrast to the linear axon found in most control corpuscles, in which branching is restricted to the ultraterminal region of the axon (Figure 1B). In some cases, the abnormal branches present in Pacinian corpuscles of *Cdx2-Cre Piezo2 cKO* mutants formed additional ultraterminal regions (Supplemental Figure 1C).

In mouse trunk hairy skin, three subsets of LTMRs form lanceolate endings, thin axonal projections that wrap around the base of hair follicles and contribute to the detection of hair movement and indentation of nearby skin: C-LTMRs, Aδ-LTMRs, and Aβ rapidly adapting (RA)-LTMRs (C, Aδ and Aβ refer to their conduction velocities, which are slow, intermediate, and fast, respectively) ^9,12^. In the skin of control animals, all three classes can be labeled by staining for the tubulin protein TUBB3 (TUJ1 antibody), whereas only Aβ RA-LTMR lanceolate endings exhibit NFH labeling. When examining the innervation pattern of hairy skin in *Cdx2-Cre Piezo2 cKO* animals, we were surprised to observe many hair follicles that completely lacked TUJ1^+^ lanceolate endings, revealing a profound deficit in end organ structure (Figure 1C). To better appreciate the relative contributions of the NFH^+^ Aβ-LTMRs and the NFH^-^ Aδ/C-LTMRs to this structural phenotype, the densities of hair follicles innervated by TUJ1^+^ and NFH^+^ lanceolate endings were quantified. We detected a ∼50% reduction in the percentage of hair follicles that contained TUJ1^+^ lanceolate endings in *Cdx2-Cre Piezo2 cKO* mutants, but no change in the percentage of hair follicles that contained NFH^+^ lanceolate endings, indicating a defect in the density or branching structure of Aδ-LTMRs and C-LTMRs (Figure 1E). Despite the normal density of hair follicles receiving NFH^+^ Aβ RA-LTMR innervation, close examination of the morphology of NFH^+^ lanceolate endings revealed defects in their branching pattern and organization: Fewer NFH^+^ lanceolate endings formed around hair follicles in the mutants, and those that were present displayed abnormal branching, turning along the circumferential axis in a manner rarely observed in controls (Figure 1D,F).

We next examined the peripheral morphology of other mechanosensory end organ structures including Merkel cell-innervating NFH^+^ neurons, NFH^+^ circumferential endings, and CGRP^+^ circumferential endings, which in mouse trunk hairy skin are normally formed by Aβ slowly adapting (SA)-LTMRs, Aβ field-LTMRs, and Aδ high threshold mechanoreceptors (HTMRs), respectively ^14,15,39,40^. We detected no major changes in these end organ structures in *Cdx2-Cre Piezo2 cKO* mutants, although the ratio of NFH^+^ axonal contacts per Merkel cell at touch domes around guard hair follicles was decreased compared to controls (Supplemental Figure 1D-H). In summary, several LTMR classes display abnormal mechanosensory end organ morphologies in *Cdx2-Cre Piezo2 cKO* mice, whereas CGRP^+^ Aδ-HTMRs and NFH^+^ Aβ field-LTMRs that form circumferential endings in hairy skin show no obvious change in their terminal morphologies.

Changes in the structures of mechanosensory end organs could reflect a developmental defect, as *Cdx2-Cre* turns on embryonically ^41^. To determine if Piezo2 influences the establishment of mechanosensory end organs, we examined peripheral sensory neuron morphologies at early postnatal ages. For these experiments, we focused on the development of the Pacinian and Meissner corpuscles, because of their stereotyped morphologies and the ease of analyzing these structures in young pups. Previous studies have shown that small Pacinian corpuscles are present around the rat fibula as early as postnatal day 0 (P0), and that they increase in size over development, reaching a mature morphology by P15 ^42^. We dissected the interosseous membrane around the fibulas of *Cdx2-Cre Piezo2 cKO* and control animals at P3-P5 and detected aberrant branching of Pacinian corpuscle afferents in the mutants, although branches in the mutant pups were smaller and less complex than those seen in adult mutants (Figure 2A-B). As development progressed, the expressivity of the branching phenotype increased, and by P15 it resembled that of the adult mutants (Figure 1B). In addition, we noticed that the corpuscles of *Cdx2-Cre; Piezo2 cKO* mutant animals were smaller than those of age-matched control littermates, and this difference in corpuscle sizes persisted into adulthood (Supplemental Figure 2B-C).

**Figure 2:**
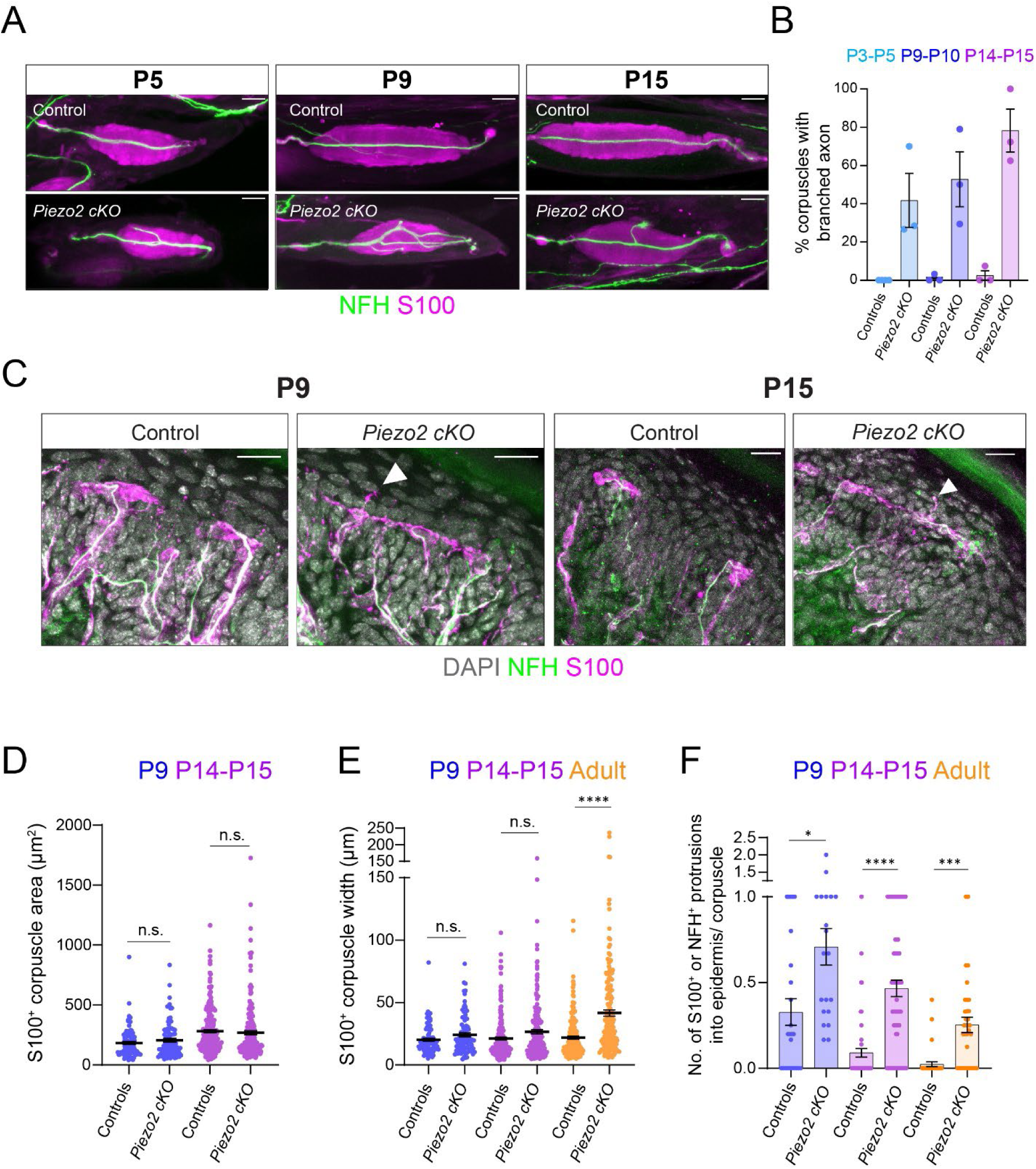
Piezo2 acts developmentally to establish mechanosensory corpuscle morphologies. **A.** Images of Pacinian corpuscles from control littermates or *Cdx2-Cre; Piezo2 cKO* mutants at postnatal days 5, 9, or 15. Tissue was stained with antibodies against NFH (green) and S100 (magenta). Scale bar = 20 µm. **B.** Quantification of the percentage of Pacinian corpuscles with branched axons at different postnatal ages. Each data point indicates one animal; averages are plotted; error bars indicate the s.e.m. See also Figure S2. **C.** Images of Meissner corpuscles in hind paw plantar pads of control littermates or *Cdx2-Cre; Piezo2 cKO* mutants at postnatal day 9 or 15. Skin tissue was stained with antibodies against NFH (green) and S100 (magenta). DAPI staining is shown in gray. Arrowheads indicate NFH^+^/S100^+^ protrusions emanating from the corpuscles and penetrating the epidermis. Scale bar = 20 µm. **D.** Quantification of the S100^+^ areas of Meissner corpuscles in control animals or *Cdx2-Cre; Piezo2 cKO* animals at different postnatal ages. There were no significant differences between areas of corpuscles in control versus mutant animals at either P9 or P14-P15 (Kruskal-Wallis test with post-hoc Dunn test for multiple comparisons). Each data point indicates one Meissner corpuscle; averages are shown by horizontal bars; error bars indicate the s.e.m. P9 data were collected from 3 controls and 3 *Cdx2-Cre; Piezo2 cKO* mutants. P14-P14 data were collected from 6 controls and 4 *Cdx2-Cre; Piezo2 cKO* mutants. **E.** Quantification of the widths of S100^+^ Meissner corpuscles at different postnatal ages. There were no differences detected between areas of corpuscles in control versus mutant animals at either P9 or P14-P15, but corpuscles in adult (>P40) animals were significantly different (****p<0.0001, Kruskal-Wallis test with post-hoc Dunn test for multiple comparisons). Each data point indicates one Meissner corpuscle; averages are shown by horizontal bars; error bars indicate the s.e.m. P9 and P14-P15 data were collected from the same animals as in 2D. Adult data were collected from the same animals as in Figure 1A. **F.** Quantification of the number of S100^+^ or NFH^+^ protrusions from Meissner corpuscles projecting into the epidermis, divided by the number of corpuscles detected in each image. *Cdx2-Cre; Piezo2 cKO* mutants formed more protrusions than control littermates at P9, P14-P15, and in adults (>P40)(*p<0.05, ***p<0.001, ****p<0.0001, adjusted p values, Kruskal-Wallis test with post-hoc Dunn test). Each point indicates one tissue section image; averages are shown by horizontal bars; error bars indicate the s.e.m. P9 and P14-P15 data were collected from the same animals as in 2D-E; adult data are collected from the same animals as in Figure 1A.

We next examined the development of Meissner corpuscles. In wild type animals, Meissner corpuscles acquire their characteristic bulbous morphologies, composed of one or two NFH^+^ axons tightly wrapped by processes from S100^+^ lamellar cells, around P9 ^43^. Corpuscles were present in hind paw glabrous skin of *Cdx2-Cre Piezo2 cKO* mutants at this age and although their morphologies appeared slightly more variable than those of controls, there was no difference in their areas or widths, suggesting that Piezo2 is not required for the initial establishment of these structures (Figure 2C-E). We noticed that many NFH^+^ or S100^+^ protrusions emerged from developing Meissner corpuscles and projected into the epidermis at P9 (Figure 2C, F). These protrusions were detected in mice of all genotypes but were more prevalent in mutants. During postnatal development, the frequency of NFH^+^ or S100^+^ epidermal protrusions decreased in controls, such that there were almost none detected in adults (Figure 2F). In *Piezo2* mutants, these structures also decreased in number over development, but to a lesser extent, and they continued to be more prevalent than in control animals at P14-P15 and in adults (Figure 2F). Taken together, our results suggest that the aberrant structures of mechanosensory end organs in *Piezo2* mutants reflect a difference in developmental trajectories and end organ formation during the first few weeks after birth, the time period when LTMRs acquire their mature peripheral morphologies and functional properties ^16,44^ (Supplemental Figure 2A).

### Piezo2 acts predominantly in neurons to influence mechanosensory end organ structures

The *Cdx2-Cre* transgene drives Cre-mediated recombination in the spinal cord and in DRGs below cervical level 2, as well as in many non-neuronal tissues ^41^. To determine if Piezo2 acts in somatosensory neurons to control mechanosensory end organ structures, we sought to genetically delete it in a cell-type specific manner. As peripheral sensory neuron-specific deletion of Piezo2 during development using *Advillin^Cre^* is lethal ^45^, we turned to a virus-based strategy to delete Piezo2. We injected AAV2/retro-hSynapsin-Cre into the paws of P3-P4 *Piezo2^flox/Null^; Rosa26^LSL-tdTomato/+^*animals and collected hind paw glabrous skin and lumbar-level DRGs four weeks after injection (Figure 3A). Cre-dependent reporter expression was enriched in neuronal axon projections in the skin, and while we observed some epidermal cell labeling, very few S100^+^ lamellar cells within the Meissner corpuscles were labeled (Supplemental Figure 3A). We analyzed the morphologies of Meissner corpuscles innervated by at least one tdTomato^+^ axon and devoid of tdTomato^+^/S100^+^ lamellar cells and observed a significant increase in the areas of these corpuscles compared to those of *Piezo2^flox/Null^*controls (Figure 3C-D). In addition, we observed more NFH^+^ or S100^+^ protrusions into the epidermis from Meissner corpuscles in the AAV Cre-injected animals compared to Cre-negative controls (Figure 3C,E).

**Figure 3:**
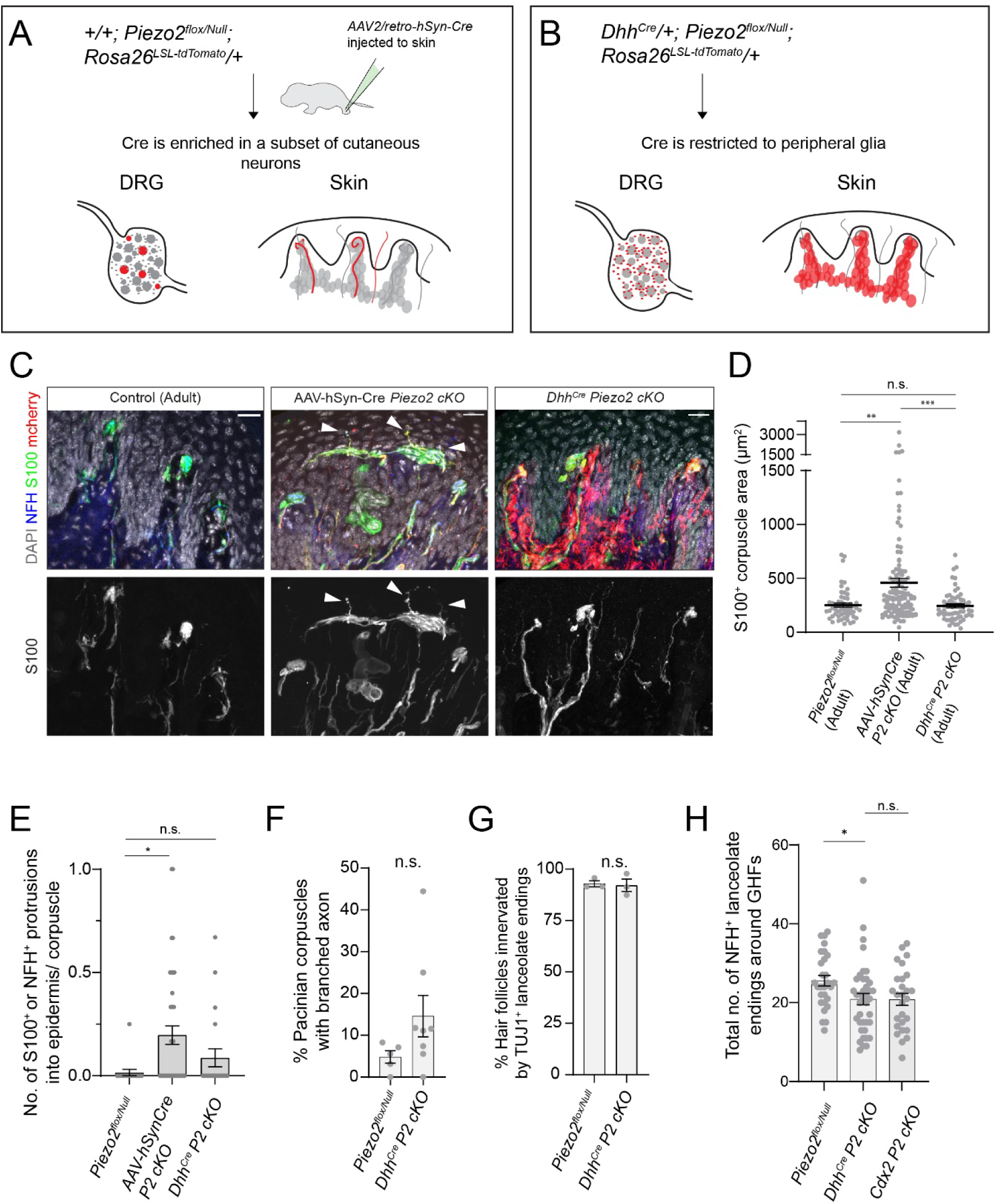
Piezo2 acts primarily in sensory neurons to influence mechanosensory end organ structures. **A.** Strategy to achieve sparse deletion of Piezo2 from neurons during early postnatal development. AAV2/retro-hSyn-Cre was injected to the hind paws of *Piezo2^flox/Null^; Rosa26^LSLtdTomato^* pups at ages postnatal day 3 to 4, and tissue was harvested at adult time points. See also Figure S3. **B.** Strategy to achieve deletion of Piezo2 from progenitors of peripheral glial cells. *Dhh^Cre/+^; Piezo2^flox/Null^; Rosa26^LSLtdTomato^* pups were bred and raised to adulthood. **C.** Images of Meissner corpuscles from a Cre-negative *Piezo2^flox/Null^* (“Control”) animal, AAV2/retro-hSyn-Cre -injected *Piezo2^flox/Null^; Rosa26^LSLtdTomato^* animal, or *Dhh^Cre/+^; Piezo2^flox/Null^; Rosa26^LSLtdTomato^* animal. Skin tissue was stained with antibodies against NFH (blue), S100 (green), and mcherry (red). DAPI staining is shown in gray. Single channel images of the S100 channel are shown below the merged color images. Arrowheads indicate S100^+^ protrusions from the corpuscles into the epidermis. Scale bar = 20 µm. **D.** Quantification of the S100^+^ areas of Meissner corpuscles in different experimental groups. AAV-hSyn-Cre injected *Piezo2^flox/Null^* animals had larger Meissner corpuscle areas than +/+; *Piezo2^flox/Null^ animals* (adjusted **p<0.001) or than *Dhh^Cre^/+; Piezo2^flox/Null^* animals (adjusted ***p<0.0001, Kruskal-Wallis test with post-hoc Dunn test for multiple comparisons). Each point indicates one Meissner corpuscle; averages are shown by horizontal bars; error bars indicate the s.e.m. Data were collected from the following animals: +/+; *Piezo2^flox/Null^*(n=3), AAV-hSyn-Cre injected +/+; *Piezo2^flox/Null^* (n=3), *Dhh^Cre/+^; Piezo2^flox/Null^* (n=4). **E.** Quantification of the number of S100^+^ or NFH^+^ protrusions from Meissner corpuscles projecting into the epidermis, divided by the number of corpuscles detected in each image. Meissner corpuscles of AAV-hSyn-Cre injected *Piezo2^flox/Null^*animals contained more S100^+^ or NFH^+^ protrusions into the epidermis than corpuscles in *Piezo2^flox/Null^ animals* (adjusted *p<0.05, Kruskal-Wallis test with post-hoc Dunn test). Each point indicates one tissue section; averages are shown by horizontal bars; error bars indicate the s.e.m. Data were collected from the same animals as in 3D. **F.** % of Pacinian corpuscles with branched axons. Each data point indicates one animal; averages are plotted; error bars indicate the s.e.m. There was no significant difference in Pacinian corpuscle branching between genotypes (Welch’s t test). Data were collected from the following animals: *+/+; Piezo2^flox/Null^* (n=5), *Dhh^Cre/+^; Piezo2^flox/Null^* (n=8). **G.** *Dhh^Cre/+^; Piezo2^flox/Null^* animals show no change in the percentage of hair follicles (HFs) innervated by TUJ1^+^ lanceolate endings compared to controls (Mann-Whitney U test). Each data point indicates one animal; averages are plotted; error bars indicate the s.e.m. Data were collected from the following animals: *+/+; Piezo2^flox/Null^* (n=3), *Dhh^Cre/+^; Piezo2^flox/Null^*(n=3). **H.** Quantification of NFH^+^ axonal structures formed by lanceolate endings around guard hair follicles. *DhhCre/+; Piezo2^flox/Null^* animals formed fewer NFH^+^ lanceolate endings compared to *Piezo2^flox/Null^* controls (*adjusted p<0.05, Kruskal-Wallis test with post-hoc Dunn test). Each data point indicates one hair follicle; averages are shown by horizontal bars; error bars indicate the s.e.m. Data were collected from the following animals: *+/+; Piezo2^flox/Null^*(n=5), *Dhh^Cre^/+; Piezo2^flox/Null^* (n=5), *Cdx2-Cre/+; Piezo2^flox/flox^*(n=3) and *Cdx2-Cre/+; Piezo2^flox/Null^* (n=2). *Cdx2-Cre; cKO* data are re-plotted from Figure 1E.

These phenotypes are similar to those observed in *Cdx2-Cre Piezo2 cKO* animals, although not as strong (Supplemental Figure 3B). Importantly, *in situ* hybridization experiments revealed that ∼30% of *tdTomato*^+^/*Nefh^+^* or *Cre^+^/Nefh*^+^ cells in lumbar-level DRGs of AAV-injected mutants retained expression of the floxed *Piezo2* exons (Supplemental Figure 3C). In contrast, *Cdx2-Cre* mediated recombination of *Piezo2^flox^* leads to almost no detectable *Piezo2*^+/^*Nefh*^+^ cells in lumbar-level DRGs ^32^. Therefore, though viral-mediated delivery of Cre leads to incomplete deletion of *Piezo2* in neurons, this manipulation partially recapitulates the effect of *Cdx2-Cre*-mediated *Piezo2* deletion on Meissner corpuscle morphology, revealing a neuronal requirement for Piezo2 in the development of these structures.

Our recent analysis of Piezo2 expression and localization found that endogenously produced Piezo2-tagged protein is enriched in the neuronal projections of LTMR axons that innervate Meissner corpuscles, Pacinian corpuscles, and hair follicles, and is not detected in the associated non-neuronal cells in these structures ^33^. However, protein levels may be below the detection threshold of immunohistochemistry, and Piezo2 may be expressed and function in non-neuronal cells during development. To test whether Piezo2 also acts in peripheral glial cells to influence mechanosensory end organ formation, we used *Dhh^Cre^*-mediated deletion. This line drives Cre expression in Schwann cell precursors, including those that develop into satellite glia in the DRG, as well as the lamellar cells and terminal Schwann cells (TSCs) that interact with mechanosensory axons in the periphery ^46^ (Figure 3B, Supplemental Figure 3D).

We first analyzed the morphologies of Meissner corpuscles following glial-specific deletion of Piezo2. A Cre-dependent reporter (*Rosa26^LSL-syt-tdTomato^*) confirmed that *Dhh^Cre^* drives efficient recombination in peripheral glial cells in the paw dermis, including S100^+^ lamellar cells in Meissner corpuscles (Figure 3C). Meissner corpuscles displayed no significant change in their morphologies in *Dhh^Cre/+^;Piezo2^flox/Null^ cKO* animals (hereafter referred to as *Dhh^Cre^; Piezo2 cKO*) compared to control animals, and had significantly smaller corpuscle areas compared to AAV-hSyn-Cre injected animals, suggesting that in contrast to neuronal Piezo2, glial cell-expressed Piezo2 is not required for the structural development of the Meissner corpuscle (Figure 3C-E, Supplemental Figure 3B).

We next asked if glial cell expression of Piezo2 regulates the morphologies of other LTMR populations. We observed no significant increase in Pacinian axon branching in *Dhh^Cre^; Piezo2 cKO* conditional mutants when compared to Cre-negative *Piezo2^flox/Null^* controls (Figure 3F), and no change in the areas or lengths of Pacinian corpuscles (Supplemental Figure 3E-F), in contrast to the effects of *Cdx2-Cre* mediated deletion. Similarly, we measured the innervation density of TUJ1^+^ lanceolate endings in trunk hairy skin and found no change in *Dhh^Cre^; Piezo2 cKO* animals, suggesting that C-LTMRs and Aδ-LTMRs innervate the skin in a normal pattern in these mutants (Figure 3G). Interestingly, when the morphology of NFH^+^ lanceolate endings was analyzed, an intermediate phenotype was observed: Fewer NFH^+^ lanceolate endings formed around guard hair follicles when compared to controls, and this phenotype was similar between *Dhh^Cre^ Piezo2 cKO* mutants and *Cdx2-Cre Piezo2 cKO* mutants (Figure 3H). In contrast, there was no significant increase in the number of turns formed by NFH^+^ lanceolate endings around hair follicles in *Dhh^Cre^ Piezo2 cKO* mutants, unlike what is observed in *Cdx2-Cre Piezo2 cKO* mutants (Supplemental Figure 3G). At non-guard hair follicles, NFH^+^ lanceolate endings were not affected by *Dhh^Cre^ -* mediated deletion of Piezo2 (Supplemental Figure 3G-H). Together, these results suggest that while Piezo2 functions primarily in neurons to establish mechanosensory end organ structures, it also plays a minor role in the glial cells of hairy skin to shape the structural development of NFH^+^ lanceolate endings. In light of these findings, and because electrophysiology experiments in *Dhh^Cre^; Piezo2 cKO* animals revealed no deficits in functional responses of Aβ-LTMRs to mechanical stimulation ^33^, we focused on the role of neuronal Piezo2 for the remainder of this study.

### Piezo2 is required for the organization of somatosensory neuron central projections

In the visual system, neural activity plays an important role in refining the topographic organization and synaptic outputs of primary sensory neuron axonal projections to central targets^47,48^. Therefore, we asked if the loss of Piezo2 impacts the central projections of somatosensory neurons. All LTMRs innervate the spinal cord dorsal horn, and a subset (Aβ-LTMRs) also forms an axonal branch that ascends the dorsal column and innervates the brainstem. To determine if this ascending projection is present in *Cdx2-Cre Piezo2 cKO* mutants, we sparsely labeled Aβ-LTMR central projections by injecting AAV-syn-tdTomato or AAV-GFP into hairy or glabrous hind paw skin and stained for fluorescently labeled presynaptic puncta in the brainstem. In all mutants with sparsely labeled neurons, Aβ-LTMRs were found to ascend the dorsal column, innervate the gracile nucleus, and form VGLUT1^+^ boutons there, consistent with our previous findings that paw-innervating Aβ-LTMRs in *Cdx2-Cre Piezo2 cKO* mutants form functional synapses with neurons in the brainstem ^32^. There was no detectable change in the overall distribution of VGLUT1^+^ signal in the gracile and cuneate nuclei in *Cdx2-Cre Piezo2 cKO* mutants, or in the co-localization of VGLUT1^+^ signal with fluorescently labeled Aβ-LTMR presynaptic terminals, when compared to wild type animals (Supplemental Figure 4A).

We next assessed synapse formation and laminar organization in the spinal cord by staining tissue sections with a panel of antibodies to detect presynaptic boutons formed by LTMRs, proprioceptors, and corticospinal neurons (VGLUT1^+^), as well as central projections formed by peptidergic (CGRP^+^) or nonpeptidergic (IB4^+^) thermoreceptors, itch receptors, and high threshold mechanoreceptors. *Cdx2-Cre Piezo2 cKO* mutants have a large reduction in VGLUT1^+^ puncta in the dorsal horn of the spinal cord, especially in lamina III-IV, the area normally targeted by LTMR axons (the LTMR-recipient zone) ^49^ (Figure 4A-B). In contrast, the projections of CGRP^+^ and IB4^+^ somatosensory neurons, normally largely restricted to lamina I-II, expanded ventrally in the mutants, encroaching into lamina III-IV (Figure 4A,C,D).

**Figure 4:**
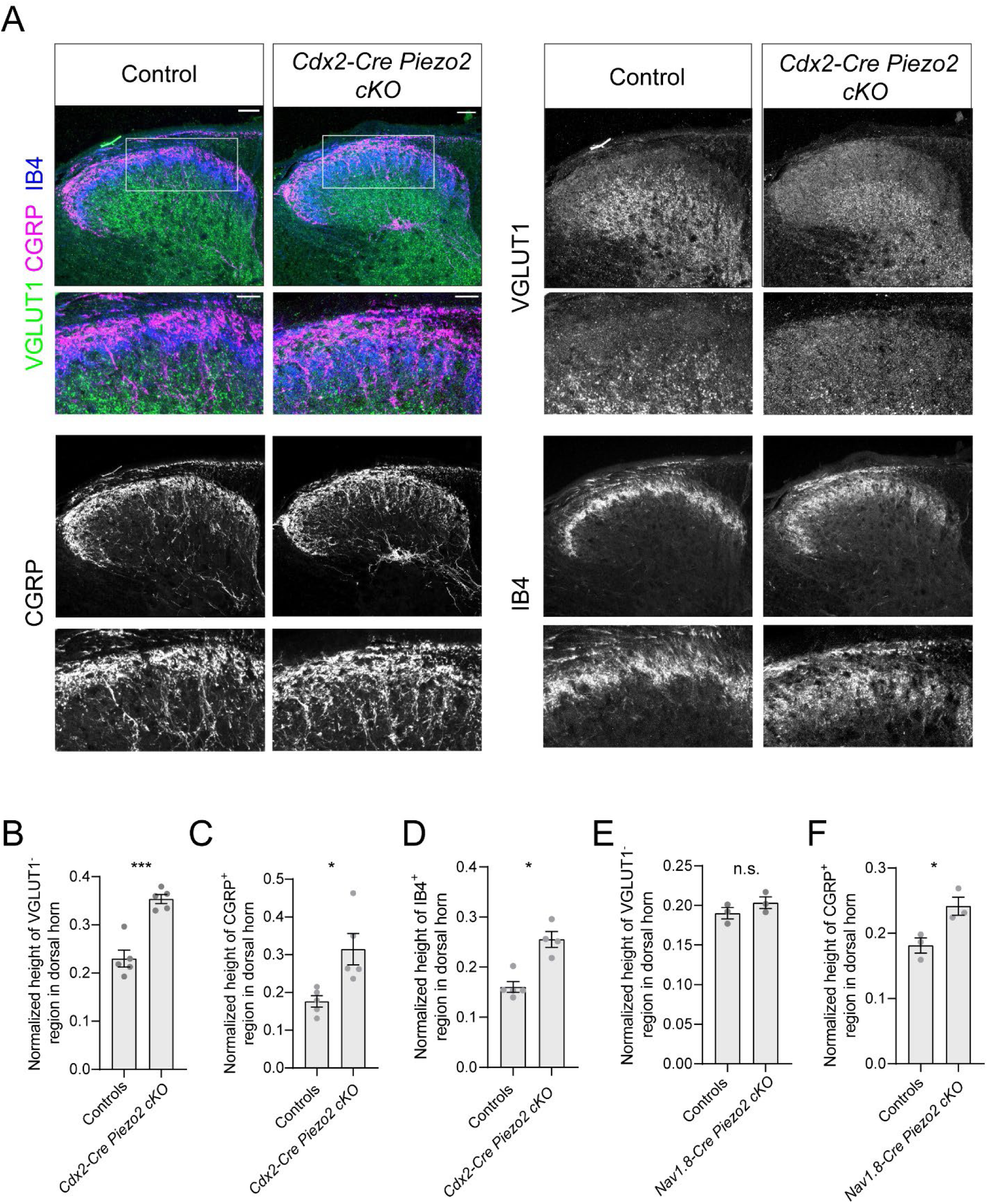
Piezo2 is required for the organization of somatosensory neuron projections in the spinal cord. **A.** Example images of thoracic-level spinal cord sections from control or *Cdx2-Cre; Piezo2 cKO* animals. One half of the dorsal horn is shown; insets show enlarged images of the areas outlined by the rectangles. Sections were stained using antibodies against VGLUT1 (green), CGRP (magenta), and for IB4-binding (blue). In *Cdx2-Cre Piezo2 cKO* mutants, the VGLUT1-negative region of the dorsal horn extends deeper into the spinal cord. In contrast, more CGRP^+^ and IB4^+^ afferents project outside of the superficial laminae. Scale bar = 50 µm in top images, 25 µm in enlarged images. **B.** Quantification of the height of the VGLUT1-negative region in the dorsal horn of the spinal cord, normalized to the height of the dorsal horn. In *Cdx2-Cre; Piezo2 cKO* mutants, the VGLUT1-negative region was larger than in control littermates (***p<0.001, unpaired t-test). Each data point indicates one animal; averages are shown by horizontal bars; error bars indicate the s.e.m. **C.** Quantification of the height of the CGRP^+^ region in the dorsal horn of the spinal cord, normalized to the height of the dorsal horn. In *Cdx2-Cre; Piezo2 cKO* mutants, the CGRP^+^ region was larger than in control littermates (*p<0.05, unpaired t-test). Each data point indicates one animal; averages are shown by horizontal bars; error bars indicate the s.e.m. **D.** Quantification of the height of the IB4^+^ region in the dorsal horn of the spinal cord, normalized to the height of the dorsal horn. In *Cdx2-Cre; Piezo2 cKO* mutants, the IB4^+^ region was larger than in control littermates (*p<0.05, Mann-Whitney U test). Each data point indicates one animal; averages are shown by horizontal bars; error bars indicate the s.e.m. **E.** Quantification of the height of the VGLUT1-negative region in the dorsal horn of the spinal cord, normalized to the height of the dorsal horn. In *Na_v_1.8-Cre; Piezo2 cKO* mutants, the VGLUT1-negative region was not different than in control littermates (unpaired t-test). Each data point indicates one animal; averages are shown by horizontal bars; error bars indicate the s.e.m. See also Figure S4. **F.** Quantification of the height of the CGRP^+^ region in the dorsal horn of the spinal cord, normalized to the height of the dorsal horn. In *Na_v_1.8-Cre; Piezo2 cKO* mutants, the CGRP^+^ region was slightly expanded relative to control littermates (*p<0.05, unpaired t-test). Each data point indicates one animal; averages are shown by horizontal bars; error bars indicate the s.e.m.

To determine if the changes in the projection patterns of CGRP^+^ and IB4^+^ neurons reflect a non-cell autonomous effect of deleting Piezo2 in LTMRs, or a cell-type autonomous function for Piezo2 in these neuronal populations, we examined the spinal cords of *Na_v_1.8-Cre; Piezo2^flox/Null^ cKO* animals. In these mutants, *Piezo2* is deleted from most small and many intermediate diameter DRG neurons that terminate within the CGRP^+^ and IB4^+^ layers, but not from large diameter Aβ-LTMRs ^50,51^. Interestingly, there was no change in the distribution of VGLUT1^+^ or IB4^+^ signal in the spinal cords of *Na_v_1.8-Cre Piezo2 cKO* mutants, but the CGRP^+^ layer once again expanded ventrally, as in *Cdx2-Cre; Piezo2 cKO* animals (Figure 4E-F, Supplemental Figure 4B-C). These results reveal that Piezo2 is required in Nav1.8^+^ neurons for regulating the central projection pattern of CGRP^+^ neurons and suggest a non-cell autonomous role for Piezo2 in regulating spinal cord projections of IB4^+^ nonpeptidergic neurons. Notably, we previously showed that corticospinal innervation of the spinal cord is also disrupted non cell-autonomously in *Cdx2-Cre; Piezo2 cKO* animals ^52^, and this likely contributes to the decreased VGLUT1^+^ signal in the dorsal horn. In summary, our anatomical analyses reveal a role for Piezo2 in regulating spinal cord projections formed by multiple classes of primary somatosensory neurons.

### Mechanosensory neurons display changes in gene expression in the absence of Piezo2

The altered morphologies of somatosensory neurons prompted us to ask whether the transcriptional profiles of DRG neurons are also impacted in the absence of Piezo2. We collected DRGs from axial levels below C2 of juvenile (P21-P24) *Cdx2-Cre Piezo2 cKO* animals and control littermates and prepared cDNA libraries for droplet-based single cell sequencing using the 10x Genomics platform. All major DRG neuronal cell types were detected in both groups and were identified using previously established markers ^53-56^. Following principal component analysis, neurons largely clustered by molecular identity, and not by genotype, with one clear exception (Figure 5A-B). The clusters that expressed canonical C-LTMR markers (*Th, Slc17a8, Nxpe2, Cckar, Cd34*) contained only control cells, and a distinct cluster composed exclusively of *Piezo2 cKO* cells expressed some C-LTMR specific genes (e.g., *Cacna1i*, *Dgkk*, *Cdh9*, *Cdh10*, *Hcrtr2*) (Supplemental Figure 5A). These neurons could not be assigned to any other cell type, as they did not express *Calca*, *Ntrk1*, *Mrgprd*, *Mrgpra3*, *Mrgprb4*, *Sst*, *Trpm8*, or *Nefh* (Supplemental Figure 5A). Therefore, this cluster is referred to as “*Piezo2 cKO* C-LTMRs.”

**Figure 5:**
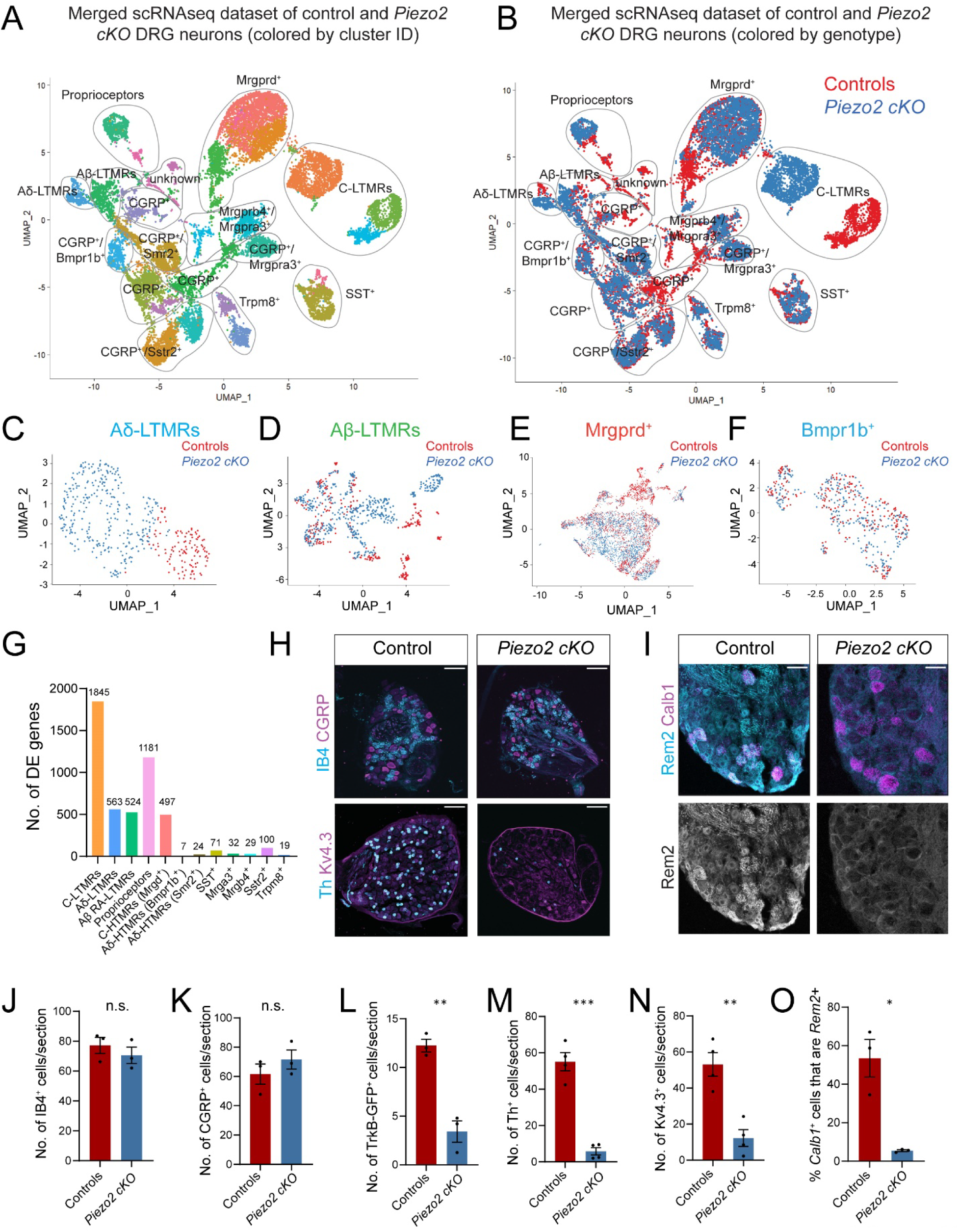
Mechanosensory neuron gene expression is altered in DRG neurons of *Piezo2 cKO* animals. **A.** UMAP plot of DRG neurons collected from 5 juvenile (P21-P24) control and 5 littermate *Cdx2-Cre; Piezo2 cKO* animals. Neurons are color-coded by computationally defined cluster identities. Molecular classes were manually assigned based on markers defined in the literature. See also Figure S5. **B.** UMAP plot of DRG neurons collected from juvenile (P21-P24) control (red) and *Piezo2 cKO* (blue) animals. Neurons are color-coded by genotype. **C.** UMAP plot of neurons assigned as Aδ-LTMRs (Cluster 17, see Supplemental Figure 5A), which were subsetted from the main dataset and re-analyzed. Control (red) and *Piezo2 cKO* (blue) neurons cluster partly by genotype. **D.** UMAP plot of neurons assigned as Aβ-LTMRs (Cluster 10), which were subsetted from the main dataset and re-analyzed. Control (red) and *Piezo2 cKO* (blue) neurons cluster partly by genotype. **E.** UMAP plot of neurons assigned as *Mrgprd*^+^ C-HTMRs (Clusters 0, 2, 8, 25), which were subsetted from the main dataset and re-analyzed. Control (red) and *Piezo2 cKO* (blue) neurons cluster partly by genotype. **F.** UMAP plot of neurons corresponding to *Bmpr1b*^+^ Aδ-HTMRs (Cluster 16), which were subsetted from the main dataset and re-analyzed. Control (red) and *Piezo2 cKO* (blue) neurons do not cluster by genotype. **G.** Number of differentially expressed (DE) genes between control and *Cdx2-Cre; Piezo2 cKO* DRG neurons of different molecularly defined classes. DE genes were identified using the Wilcoxon rank-sum test, adjusted p value <0.05, log-fold change ≥0.5. **H.** Top: Images of thoracic-level DRG sections from control or *Cdx2-Cre; Piezo2 cKO* animals stained with anti-CGRP (magenta) or IB4-conjugated (cyan) antibodies. Bottom: Images of thoracic-level DRG sections from control or *Cdx2-Cre; Piezo2 cKO* animals stained with anti-K_v_4.3 (magenta) or anti-Th (cyan) antibodies. Scale bar = 100 µm. **I.** Images of lumbar-level DRG sections from control or *Cdx2-Cre; Piezo2 cKO* animals stained with RNAscope probes to detect *Calb1* (magenta, a marker for Aβ-LTMRs) or *Rem2* (cyan) transcripts. Scale bar = 50 µm. **J.** Quantification of the number of IB4^+^ cells in thoracic DRG sections for control or *Cdx2-Cre; Piezo2 cKO* animals. There was no difference between the two groups (unpaired t-test). Each data point indicates one animal; averages are shown by horizontal bars; error bars indicate the s.e.m. **K.** Quantification of the number of CGRP^+^ cells in thoracic DRG sections for control or *Cdx2-Cre; Piezo2 cKO* animals. There was no difference between the two groups (unpaired t-test). Each data point indicates one animal; averages are shown by horizontal bars; error bars indicate the s.e.m. **L.** Quantification of the number of *TrkB^GFP+^* cells in thoracic DRG sections for control or *Cdx2-Cre; Piezo2 cKO* animals. There was a reduction in the mutants (**p<0.01, unpaired t-test). Each data point indicates one animal; averages are shown by horizontal bars; error bars indicate the s.e.m. **M.** Quantification of the number of Th*^+^* cells in thoracic DRG sections for control or *Cdx2-Cre; Piezo2 cKO* animals. There was a reduction in the mutants (***p=0.0001, unpaired t-test). Each data point indicates one animal; averages are shown by horizontal bars; error bars indicate the s.e.m. **N.** Quantification of the number of K_v_4.3*^+^* cells in thoracic DRG sections for control or *Cdx2-Cre; Piezo2 cKO* animals. There was a reduction in the mutants (**p<0.01, unpaired t-test). Each data point indicates one animal; averages are shown by horizontal bars; error bars indicate the s.e.m. **O.** Quantification of the percentage of *Calb1*^+^ cells that are *Rem2*+ in lumbar DRG sections for control or *Cdx2-Cre; Piezo2 cKO* animals. There was a reduction in the mutants (*p<0.05, Welch’s t-test). Each data point indicates one animal; averages are shown by horizontal bars; error bars indicate the s.e.m.

*Cdx2-Cre Piezo2* mutant neurons expressing genes characteristic of other molecularly defined mechanosensory neuron classes, including Aδ-LTMRs (*Nefh*^+^/*Ntrk2*^+^/*Colq*^+^/*Ntng1*^+^), Aβ-LTMRs (*Nefh*^+^/*Ntng1*^+^/*Calb1*^+^), proprioceptors (*Nefh*^+^/*Ntrk3*^+^/*Pvalb*^+^/*Runx3*^+^/*Whrn*^+^), C-HTMRs (*Mrgprd^+^*) and Aδ-HTMRs (*Calca^+^/Nefh^+^/Bmpr1b^+^*or *Calca^+^/Nefh^+^/Smr2^+^*), were readily identified and clustered together with control cells (Figure 5A-B). However, when Aδ-LTMRs were subsetted from the main dataset and re-analyzed, control and mutant neurons clustered partly by genotype, suggesting differences in gene expression (Figure 5C). Similar effects were observed with Aβ-LTMRs, proprioceptors, and *Mrgprd^+^* neurons (Figure 5D,E, Supplemental Figure 5B). In contrast, when clusters corresponding to Aδ-HTMRs (*Bmpr1b^+^* or *Smr2^+^*), pruriceptors (*SST^+^*, *Mrgpra3^+^*, or *Mrgprb4^+^*) or thermoreceptors (*Calca^+^/Sstr2^+^* or *Trpm8^+^*) ^54^ were subsetted and re-analyzed, cells did not segregate by genotype, suggesting minimal differences in gene expression between control and mutant neurons of these classes (Figure 5F, Supplemental Figure 5B).

To identify genes that are differentially expressed between *Cdx2-Cre Piezo2 cKO* and control neurons, we performed differential gene expression analysis in a subset-specific manner. Hundreds to thousands of genes were differentially expressed between mutant and control C-LTMRs, Aδ-LTMRs, Aβ-LTMRs, proprioceptors, and *Mrgprd*^+^ neurons (Figure 5G). In contrast, few differences in gene expression were detected when comparing mutant and control CGRP^+^ Aδ-HTMRs *(Bmpr1b*^+^ or *Smr2*^+^), pruriceptors (*SST^+^*, *Mrgpra3^+^*, or *Mrgprb4^+^*), warm-sensitive thermoreceptors (*Calca*^+^/*Sstr2*^+^), or cold-sensitive thermoreceptors (*Trpm8*^+^). Interestingly, genes with altered expression levels in *Piezo2 cKO* LTMRs included several that encode cell surface receptors or cytosolic signaling molecules with known roles in axon guidance or neuronal morphogenesis (*Nrp2*, *Rem2*, *Epha4*, *Sema5a*) ^57-63^, suggesting possible mechanisms by which Piezo2 might indirectly regulate axonal development or outgrowth (Supplemental Figure 5G-I).

To validate our sequencing results, we first examined the expression of common markers for DRG cell types by performing immunohistochemistry on tissue sections. We detected no change in the expression patterns of NF200, CGRP, or in IB4 binding in *Cdx2-Cre Piezo2 cKO* mutants, consistent with the normal expression of *Nefh*, *Calca*, and markers of non-peptidergic DRG neurons as detected by RNA sequencing (Figure 5H, J, K, Supplemental Figure 5E). In contrast, fewer *TrkB^tauEGFP+^*, Th^+^, and K_v_4.3^+^ cells were detected by immunohistochemistry in *Cdx2-Cre Piezo2 cKO* DRGs, supporting the decreased expression of *Ntrk2*, *Th*, and *Kcnd3* RNA in our bioinformatic analyses (Figure 5H, L-M, Supplemental Figure 5F).

Next, we performed *in situ* hybridization (RNAscope) experiments to assess expression of *Th*, *Nrp2,* an*d Rem2*, which were differentially expressed in C-LTMRs, Aδ-LTMRs, or Aβ-LTMRs, respectively. We observed strong reductions in the signal for all three probes in DRGs of *Cdx2-Cre Piezo2 cKO* mutants (Figure 5I, O and Supplemental Figure 5C-D). *Rem2,* the most highly differentially expressed gene in *Cdx2-Cre Piezo2 cKO* Aβ-LTMRs, encodes a Ras-like GTPase protein that regulates neuronal morphology, synapse formation, and neuronal excitability in visual circuits, where its expression is activity-dependent ^62-65^. Impressively, almost no *Rem2*^+^ cells were detected in lumbar-level DRGs of *Cdx2-Cre Piezo2 cKO* animals (Figure 5I). To determine if Rem2 plays a role in establishing LTMR peripheral morphologies, we collected peripheral tissues from global *Rem2* knock-out (null) animals. Pacinian and Meissner corpuscles had normal morphologies in the homozygous mutants, and there was no difference in the total number of NFH^+^ lanceolate endings around hair follicles (Supplemental Figure 5J-M). However, we observed a small increase in the average number of turns formed by lanceolate endings around guard hair follicles in *Rem2* mutants, although this phenotype is considerably weaker than that observed in *Cdx2-Cre Piezo2 cKO* mutants (Supplemental Figure 5N). Thus, although Rem2 is not a primary downstream effector of Piezo2 in controlling mechanosensory end organ morphogenesis, it may help fine-tune the morphology of a subset of Aβ-LTMRs that innervate hairy skin. Together, these findings show that the general transcriptional landscape of DRG neurons is preserved in the absence of Piezo2, but that gene expression patterns in sensory neuron populations normally activated by light mechanical forces are significantly altered, with a particularly strong effect on the differentiation of C-LTMRs.

### The voltage-gated sodium channel Na_v_1.6 is required for high frequency responses to tactile stimuli and for normal peripheral mechanosensory neuron morphologies

How does Piezo2 regulate mechanosensory neuron structural maturation? One possibility is that Piezo2 is required for stimulus-dependent neuronal activity, which in turn controls LTMR morphogenesis and end organ formation, through downstream signaling cascades that impinge on gene regulation or cytoskeletal remodeling. Alternatively, Piezo2 could be acting independently of its ion channel function. Indeed, a recent study revealed a role for *Drosophila* Piezo during olfactory projection neuron dendrite targeting that does not require its channel activity ^66^. To distinguish between these two models, we sought to manipulate neural activity by an alternate method and assess the consequences on maturation of LTMR peripheral end organs. For these experiments, we focused on A fiber-LTMRs because they are easily accessible for *in vivo* recordings and form end organ structures that are highly disorganized in *Piezo2* mutants (Meissner and Pacinian corpuscles, and the lanceolate endings around hair follicles). Of the nine voltage-gated sodium channels in mice, Na_v_1.1 (encoded by the gene *Scn1a*), Na_v_1.6 (*Scn8a*), and Na_v_1.7 (*Scn9a*) are highly expressed in Aβ-LTMRs during development and into adulthood ^53,67^. We first generated animals in which *Scn8a* (Na_v_1.6) is deleted from peripheral sensory neurons using *Advillin^Cre^*. Conditional mutants are born in Mendelian ratios and survive to adulthood, although they display ataxia by weaning ages. Consistent with this observation, a recent study reported a motor coordination phenotype in animals in which *Scn1a* (Na_v_1.1) is deleted from somatosensory neurons and proposes that Na_v_1.1 and Na_v_1.6 function together in proprioceptors to generate normal firing patterns in response to muscle stretch ^68^.

To determine if mechanically evoked activity in Aβ-LTMRs is affected in *Avil^Cre/+^; Scn8a^flox/flox^ cKO* mutants (hereafter referred to as *Avil^Cre^; Scn8a cKO*), we performed single unit *in vivo* juxtacellular recordings of DRGs in juvenile animals (P21-P30)(Figure 6A). We delivered a vibration stimulus to the skin to identify A-fiber LTMRs. In controls, applying a 200-300 Hz vibration stimulus to the hind limb skin allowed us to detect responses from three distinct classes of mechanically sensitive units: 1) Pacinian corpuscle afferents, defined here as those able to fully entrain to 200 Hz or higher frequency vibration at forces ≤ 10 mN; 2) hairy skin LTMRs, which respond best to gentle brushing or deflection of hairs but are also sensitive to skin indentation, vibration, and air puff; and 3) proprioceptors, which are most robustly activated by paw or leg movement, but can also respond to vibration. Strikingly, in *Avil^Cre^; Scn8a cKO* animals, no units with Pacinian-like responses were detected, although hairy skin LTMRs and proprioceptors were readily identified (n=7 animals) (Figure 6B, E-H, Supplemental Figure 6A). In contrast, units with Pacinian-like responses were identified in 7 out of 10 control mice (Figure 6B, G).

**Figure 6:**
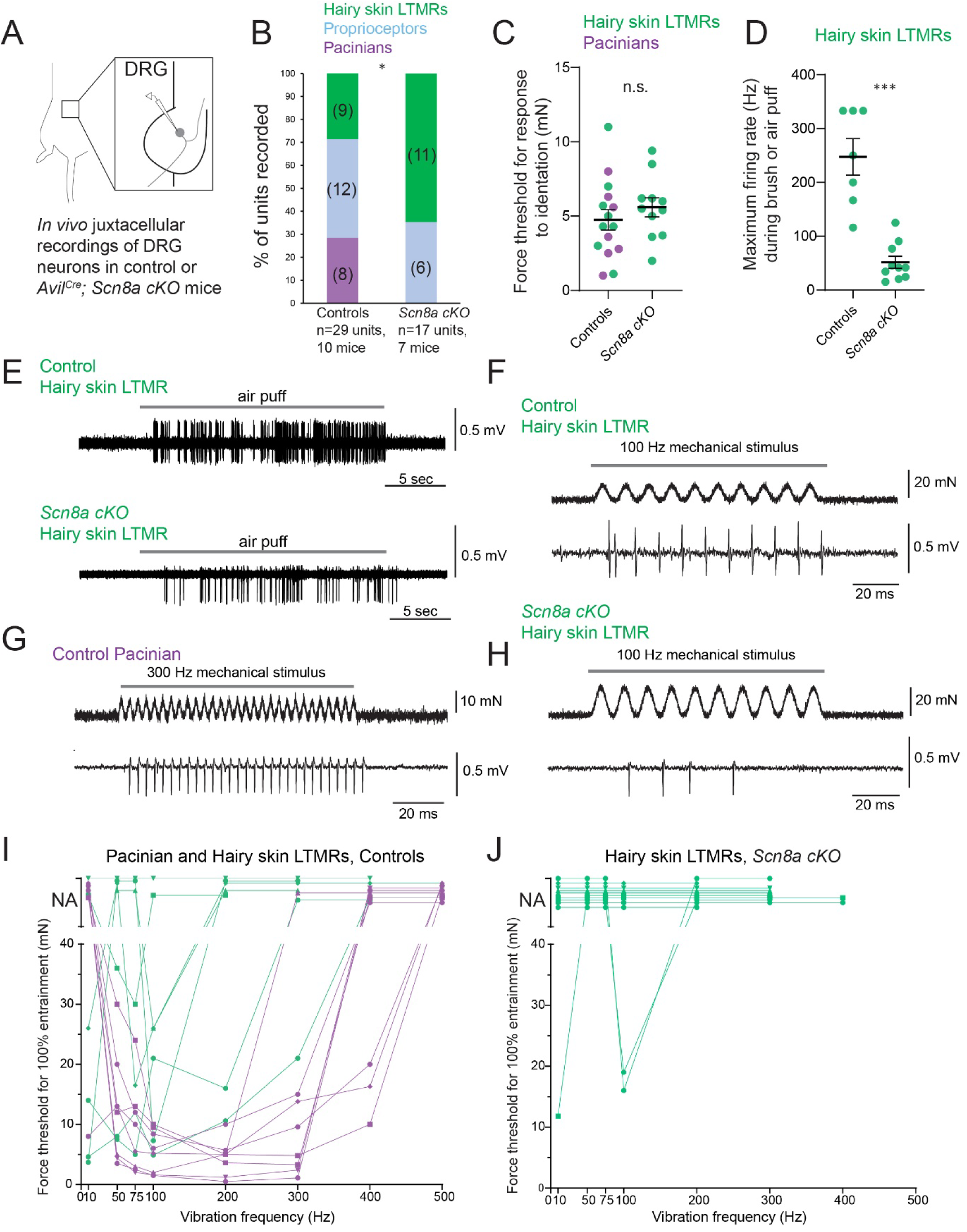
Na_v_1.6 is required for high frequency responses to mechanical stimuli in DRG neurons. **A.** Schematic of the experimental preparation. L4 (lumbar-level 4) DRGs were exposed in anesthetized mice, and single unit juxtacellular recordings were performed. **B.** Summary of the three classes of mechanosensory neurons that were detected using the mechanical search stimulus. In controls, hairy skin LTMRs (green), proprioceptors (blue), and Pacinian afferents (purple) were identified. In Na_v_1.6 (*Scn8a*) cKO animals, hairy skin LTMRs (green) and proprioceptors (blue) were detected (*p<0.05, Fisher’s exact test). Data were collected from 10 control animals and 7 *Avil^Cre/+^; Scn8a^flox/flox^* animals. See also Figure S6. **C.** The minimum force required for a unit to be activated by indentation. There was no difference between LTMRs recorded from controls or from *Scn8a cKO* mutants (unpaired t test). Each data point indicates one neuron; averages are shown by horizontal bars; error bars indicate the s.e.m. **D.** The instantaneous maximum firing rate was obtained for hairy skin LTMRs during 20 second stimulus intervals when neurons were mechanically activated by gentle brush of the skin or by air puff. Maximum firing rates were reduced in *Scn8a cKO* mutants (***p<0.001, Welch’s t test). Each data point indicates one neuron; averages are shown by horizontal bars; error bars indicate the s.e.m. **E.** Top: *In vivo* recording of a hairy skin LTMR from a control animal (*+/+; Scn8a^flox/+^*). The gray rectangle indicates the time window during which air puff was delivered to the unit’s receptive field, which was on the thigh. Bottom: *In vivo* recording of a hairy skin LTMR from an *Avil^Cre/+^; Scn8a^flox/flox^*animal. The gray rectangle indicates the time window during which air puff was delivered to the unit’s receptive field, which was on the thigh. **F.** *In vivo* recording of a hairy skin LTMR from a control animal (*+/+; Scn8a^flox/+^*, same unit as shown in Figure 6E). A 100 Hz vibration stimulus was delivered to the unit’s receptive field, which was on the thigh. **G.** *In vivo* recording of a Pacinian unit from a control animal *(+/+; +/+)*. A 300 Hz vibration stimulus was delivered to the unit’s receptive field, which was above the heel. **H.** *In vivo* recording of a hairy skin LTMR from an *Avil^Cre/+^; Scn8a^flox/flox^* animal (same unit as shown in Figure 6E). A 100 Hz vibration stimulus was delivered to the unit’s receptive field, which was on the thigh. **I.** The minimum force required for units to reach 100% entrainment (i.e., firing at least once per cycle) when responding to a vibration stimulus is plotted for control animals. Pacinian units are shown in purple; hairy skin LTMRs are in green. “NA” indicates that a unit did not reach 100% entrainment at forces up to 35 mN (higher forces were not tested). Each data point indicates one unit tested at a specific frequency; points indicating the same neuron are connected by lines. **J.** The minimum force required for hairy skin LTMR units to reach 100% entrainment (i.e., firing at least once per cycle) when responding to a vibration stimulus is plotted for *Avil^Cre^ Scn8a cKO* animals. “NA” indicates that a unit did not reach 100% entrainment at forces up to 35 mN (higher forces were not tested). Each data point indicates one unit tested at a specific frequency; points indicating the same neuron are connected by lines.

We measured the sensitivity of Pacinian and hairy skin LTMRs to mechanical indentation and to vibratory stimuli. Control and *Avil^Cre^; Scn8a cKO* mechanosensory units responded similarly to indentation (Figure 6C). In both genotypes, LTMRs were very sensitive, with an average force threshold below 10 mN. These results suggest that unlike Piezo2, Na_v_1.6 is not required for LTMR activation by low force mechanical stimuli. Pacinian units identified in control animals had a U-shaped tuning curve to vibration, with peak sensitivity in response to frequencies ranging from 100-400 Hz, similar to what has been described for Pacinian afferents in mice and other species (Figure 6I; purple) ^69,70^. Hairy skin LTMRs were able to partially entrain to vibration at lower frequencies (10-100 Hz) in both control and *Avil^Cre^; Scn8a cKO* animals, although 7/10 mutant neurons in these experiments failed to reach 100% entrainment at any of the frequencies tested, suggesting a defect in repetitive firing (Figure 6F-J, Supplemental Figure 6B-C). Indeed, while hairy skin units in *Avil^Cre^; Scn8a cKO* animals robustly fired in response to air puff or to skin brush, their maximum firing rates during these periods of mechanically evoked activity were significantly lower than those of control hairy skin LTMRs (Figure 6D-E). Together, these results show that Na_v_1.6 (*Scn8a*) is required for high frequency responses to mechanical stimuli in somatosensory neurons.

We next examined the peripheral morphologies of mechanosensory neurons in *Avil^Cre^; Scn8a cKO* mutants. Pacinian corpuscles were found in the expected location and numbers in the mutants, suggesting that the lack of Pacinian-like functional responses was not caused by the absence of this end organ. However, a significant proportion of Pacinian corpuscles in *Avil^Cre^; Scn8a cKO* animals were innervated by aberrantly branching axons, reminiscent of the morphological phenotypes observed in *Cdx2-Cre Piezo2 cKO* mutants (Figure 7A). There was no change in the areas or lengths of Pacinian corpuscles in *Avil^Cre^; Scn8a cKO* mutants, though we observed several instances of denervated corpuscles (lacking an NFH^+^ axon) in the mutants, which were almost never seen in controls (Supplemental Figure 7A-D). Impressively, Meissner corpuscles also had abnormal morphologies in *Avil^Cre^; Scn8a cKO* mutants, displaying larger areas and greater widths than in controls (Figure 7B-D). In contrast, we detected no major defects in the NFH^+^ Aβ-LTMRs that innervate hairy skin, although NFH^+^ lanceolate endings around non-guard hair follicles were slightly more numerous in mutants than in controls (Figure 7E, Supplemental Figure 7F). These results demonstrate that Na_v_1.6 is required for the normal morphologies of Aβ RA-LTMRs that innervate Meissner and Pacinian corpuscles.

**Figure 7:**
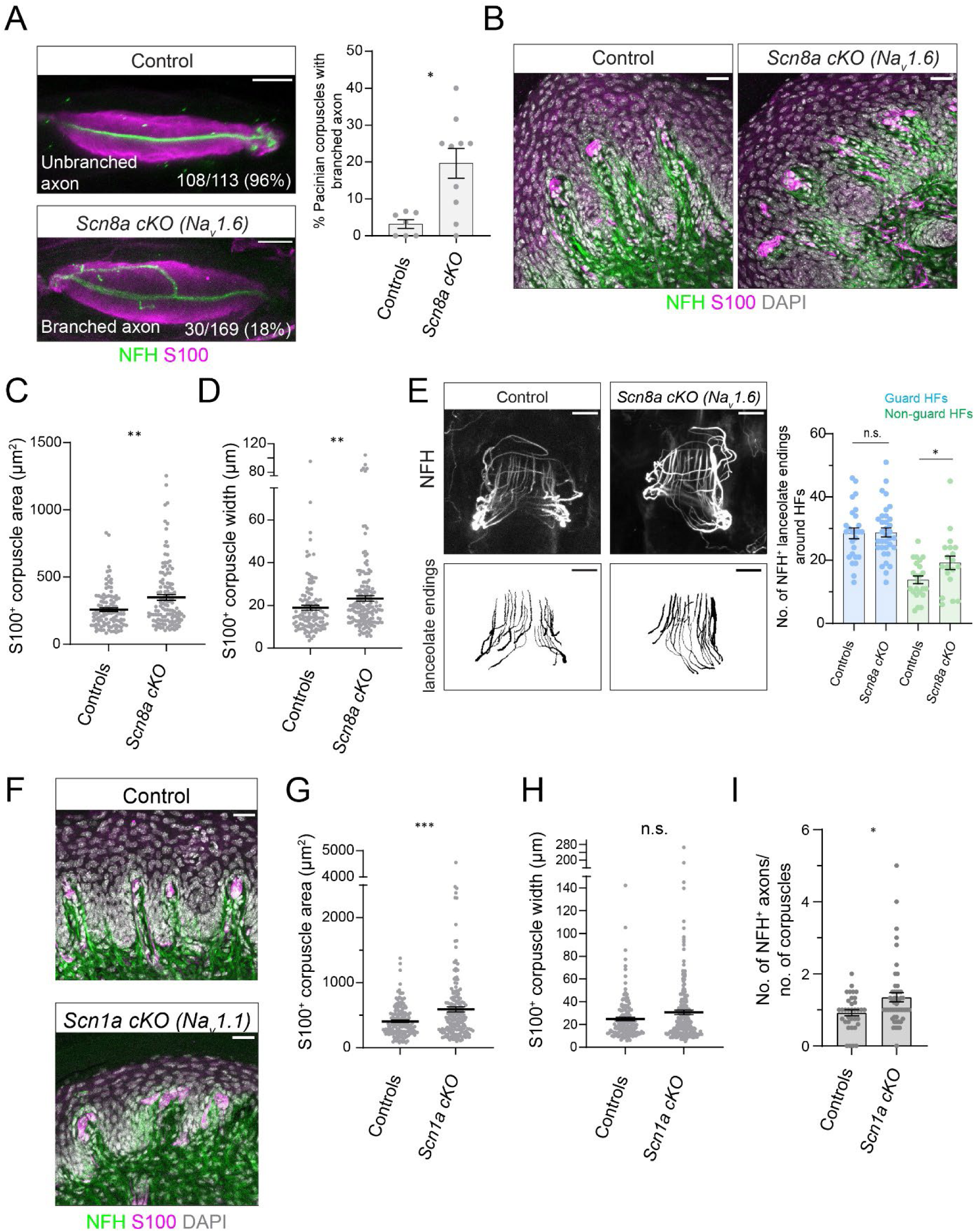
Na_v_ channels are required for normal mechanosensory end organ structures. **A.** Left: Images of Pacinian corpuscles from an adult control littermate or adult *Avil^Cre^/+; Scn8a^flox/flox^ cKO* animal. Tissue was stained using antibodies against NFH (green) and S100 (magenta). Scale bar = 25 µm. Right: Quantification of the percentage of Pacinian corpuscles with branched axons in adults (*p=0.01, Mann-Whitney U test). Each data point indicates one animal; averages are plotted; error bars indicate the s.e.m. See also Figure S7. **B.** Images of Meissner corpuscles from an adult control littermate or from an *Avil^Cre^/+; Scn8a^flox/flox^ cKO* animal. Skin tissue was stained using antibodies against NFH (green) and S100 (magenta). DAPI staining is shown in gray. Scale bar = 25 µm. **C.** Quantification of the S100^+^ areas of Meissner corpuscles in controls or *Avil^Cre/+^; Scn8a^flox/flox^ cKO* adult animals. Meissner corpuscle areas were larger in *Scn8a cKO* mutants (**p<0.01, Mann-Whitney U test). Each data point indicates one Meissner corpuscle; averages are shown by horizontal bars; error bars indicate the s.e.m. Data were collected from 5 control littermates and 5 *AvilCre/+; Scn8a^flox/flox^* mutants. **D.** Quantification of Meissner corpuscle widths in controls or *Avil^Cre/+^; Scn8a^flox/flox^ cKO* adult animals. Meissner corpuscle widths were larger in *Scn8a cKO* mutants (**p<0.01, Mann-Whitney U test). Each data point indicates one Meissner corpuscle; averages are shown by horizontal bars; error bars indicate the s.e.m. Data were collected from the same animals as in Figure 7C. **E.** Left, top: Images of guard hair follicles innervated by NFH^+^ lanceolate endings. Bottom: Reconstructions of lanceolate endings. Scale bar = 20 µm. Right: Quantification of NFH^+^ axonal structures formed by lanceolate endings around guard hairs (blue) or non-guard hair follicles (green). There was no difference in the total number of NFH^+^ lanceolate endings formed in *Scn8a cKO* animals around guard hair follicles (unpaired t-test), and an increase in the number of endings formed in the mutants around non-guard hair follicles (*p<0.05, Welch’s t test). Each data point indicates one hair follicle; averages are plotted; error bars indicate the s.e.m. Data were collected from 4 control animals and 4 *AvilCre/+; Scn8a^flox/flox^*mutants. **F.** Images of Meissner corpuscles from an adult control littermate or from an *Avil^Cre^/+; Scn1a^flox/flox^ cKO* animal. Skin tissue was stained using antibodies against NFH (green) and S100 (magenta). DAPI staining is shown in gray. Scale bar = 25 µm. **G.** Quantification of the S100^+^ areas of Meissner corpuscles in controls or *Avil^Cre/+^; Scn1a^flox/flox^ cKO* adult animals. Meissner corpuscle areas were larger in *Scn1a cKO* mutants (***p<0.001, Mann-Whitney U test). Each data point indicates one Meissner corpuscle; averages are shown by horizontal bars; error bars indicate the s.e.m. Data were collected from 5 control littermates and 6 *Avil^Cre/+^; Scn1a^flox/flox^* mutants. **H.** Quantification of Meissner corpuscle widths in controls or *Avil^Cre/+^; Scn1a^fl/fl^ cKO* adult animals. Meissner corpuscle widths were not significantly different between the two groups (Mann-Whitney U test). Each data point indicates one Meissner corpuscle; averages are shown by horizontal bars; error bars indicate the s.e.m. Data were collected from the same animals as in Figure 7G. **I.** Quantification of the number of NFH^+^ axons innervating Meissner corpuscles, normalized by the number of corpuscles in each image. Meissner corpuscles in *Avil^Cre/+^; Scn1a^flox/flox^ cKO* animals were innervated by more NFH^+^ axons than corpuscles in controls (*p<0.05, Mann-Whitney U test). Each point indicates one image; averages are shown by horizontal bars; error bars indicate the s.e.m. Data were collected from the same animals as in Figure 7G-H.

The anatomical and functional phenotypes of *Avil^Cre^; Scn8a cKO* mutants partially recapitulate what is observed in *Cdx2-Cre Piezo2 cKO* mutants but are not as pronounced. Similarly, our electrophysiology experiments revealed that mechanically evoked activity in LTMRs was incompletely disrupted in *Avil^Cre^; Scn8a cKO* mutants, in contrast to the complete loss of responses to low force mechanical stimuli observed in *Cdx2-Cre Piezo2 cKO* animals. Redundancy among the multiple Na_v_ channels expressed in LTMRs could account for the different effects of these genetic manipulations. Interestingly, a recent study showed that Na_v_1.1, Na_v_1.6, and Na_v_1.7 are co-expressed in proprioceptors and that all three channels contribute to sodium currents measured in these neurons *in vitro* ^68^. Therefore, to determine if other voltage-gated sodium channels expressed in LTMRs influence mechanosensory end organ morphologies, we generated *Avil^Cre^; Scn1a^flox/flox^ (Na_v_1.1) cKO* animals. We detected no significant defects in the morphologies of NFH^+^ lanceolate endings around hair follicles or in Pacinian corpuscles of these mutants (Supplemental Figure 7G-I). However, *Avil^Cre^ Scn1a cKO* mutants had abnormally elongated, disorganized, and hyper-innervated Meissner corpuscles, similar to the phenotype of *Avil^Cre^ Scn8a cKO* and *Cdx2-Cre Piezo2 cKO* animals (Figure 7F-I). Taken together, these findings support a model in which Piezo2-mediated activation of multiple, redundant, voltage-gated sodium channels acts during development to establish normal morphological and molecular properties of mechanosensory neurons.

## Discussion

In this study we uncover an essential role for the mechanotransduction channel Piezo2 and neural activity in establishing mechanosensory end organ structures and the transcriptional maturation of somatosensory neurons. We propose that during early life, mechanical forces generated either within developing tissues or from the external environment influence mechanosensory neuron maturation and help shape the formation of the exquisitely sensitive mechanosensory end organs that underlie our sense of light touch.

### Mechanotransduction regulates somatosensory neuron morphogenesis

Our finding that mechanotransduction shapes the development of mechanosensory end organs was unexpected. Prior studies using developmentally active Cre lines to delete Piezo2 and/or Piezo1 reported no overt structural alterations in proprioceptors ^24^, the vagal neurons that innervate the arterial ligament ^27^, or in DRG neurons that innervate the bladder ^71^. These neuronal populations may not rely on mechanically evoked activity for their maturation. Alternatively, Piezos could contribute to aspects of their development that were not analyzed in mutants, including gene expression or central synapse formation. While surprising, our findings are consistent with previous work demonstrating an inverse relationship between DRG neuron activity and neurite outgrowth ^72-74^, and with recent reports that mechanosensitive ion channels regulate neuronal morphology during development or regeneration ^75-77^. The morphological phenotypes we observed upon Piezo2 deletion are complex. In Meissner and Pacinian corpuscles, peripheral axons appeared to form larger arbors and to branch exuberantly. These phenotypes are consistent with a model in which mechanically evoked activity acts as a “brake” to inhibit outgrowth once axons reach their peripheral targets and are activated by mechanical stimuli. In contrast, the lanceolate endings formed by LTMRs around hair follicles were less frequently detected in *Cdx2-Cre Piezo2 cKO* mutants compared to wild type animals, suggesting that these neurons failed to innervate hair follicles at a normal density due to decreased axon outgrowth or terminal branching or because they formed endings with entirely different morphologies. Sparse labeling experiments in which peripheral and central arbors of LTMRs are resolved at the single cell level would be useful to assess how somatosensory axon outgrowth and developmental pruning are affected in *Piezo2* mutants.

What are the mechanisms by which Piezo2 influences the structural properties of somatosensory neurons? Our experiments in sodium channel mutants support a model in which diminished or abnormal action potential firing patterns in LTMRs lead to altered peripheral neuron morphologies. In Purkinje cells, Na_v_1.6 generates a large fraction of the resurgent sodium current that contributes to repetitive firing^78^. Na_v_1.6 is also required for resurgent sodium currents in large-diameter DRG neurons *in vitro*^79^, and our findings demonstrate that this channel is required for high-frequency firing in somatosensory neurons *in vivo*. We propose that in Na_v_1.6 (*Scn8a*) conditional mutants, a reduction in high frequency firing in Aβ-LTMRs underlies the structural changes in Meissner and Pacinian corpuscle development. Abnormal branching similar to what is detected in Piezo2 and Na_v_1.6 mutants has also been reported in the axons of Pacinian corpuscles following nerve injury and re-innervation in rats ^80^, potentially reflecting a relationship between decreased activity and increased Pacinian axon complexity. Intriguingly, peripheral glutamatergic signaling between neurons and glial cells has been proposed to control hair follicle innervation ^81^, and Piezo2 acts in proprioceptor axons to influence tendon morphogenesis through local vesicle release ^38^. Exploring the molecular mechanisms by which mechanotransduction and LTMR action potential firing control the development of end-organs and peripheral tissues offers an exciting future direction.

Does Piezo2 influence mechanosensory end organ development independent of its ion channel function? The effects of Na_v_1.1 or Na_v_1.6 deletion on somatosensory neuron morphologies are less strong than what is observed in *Cdx2-Cre Piezo2 cKO* mutants. This could be due to redundancy among the different sodium channels that are co-expressed in Aβ-LTMRs, or because Piezo2 acts by an additional mechanism that is independent of neuronal activity. Rescue experiments to restore wild type or channel-inactivated Piezo2 in different cell types may clarify if channel-independent developmental functions for Piezo2 exist.

### Activity-dependent transcriptional maturation of somatosensory neurons

The widespread changes in gene expression that we observed in *Cdx2-Cre Piezo2 cKO* mutants across multiple mechanosensory neuron populations suggest that mechanically evoked activity broadly influences somatosensory neuron transcriptional maturation. Importantly, no major changes in gene expression were observed in cells that do not express Piezo2 (*Trpm8^+^*and *Sst*^+^ populations), demonstrating that the phenotype we observed is specific to cells that rely on Piezo2-mediated mechanotransduction for their normal activity patterns. Indeed, the largest changes in gene expression were detected in neurons that respond to low force mechanical stimuli (LTMRs, proprioceptors, and *Mrgprd*^+^ neurons), with the strongest effect in C-LTMRs. Interestingly, recent studies using *Scn9a* (Na_v_1.7) conditional mutants report major changes in C-LTMR genome-wide expression and decreased expression of C-LTMR markers ^82,83^ that are remarkably similar to the effect we detected in *Cdx2-Cre Piezo2 cKO* animals. Taken together with the finding that Na_v_1.7 is required for normal C-LTMR excitability and mechanical sensitivity^84^, our results strongly support a role for mechanically evoked activity in the transcriptional maturation of the somatosensory neurons that respond to low force mechanical stimuli. Whether other Na_v_ channels regulate the transcriptional maturation of the cell types they are expressed in remains an open question, as does the relationship between transcriptional changes and altered neuronal morphologies, but it is noteworthy that some human patients with SCN9A mutations have decreased intra-epidermal nerve fiber density in skin biopsies ^85^. Thus, channelopathies associated with sensory neuron dysfunction might in some cases reflect anatomical or developmental deficits, though additional work will be required to elucidate the contributions of specific ion channels to the acquisition of mature somatosensory neuron identities and structures.

### Parallels and distinctions with other sensory systems

Our findings highlight shared principles in the role that sensory transduction and neuronal activity play in the maturation of peripheral sensory organs across modalities. In the auditory system, loss of function mutations in mechanotransduction channels cause changes in the morphology of stereocilia on inner hair cells ^86-88^, and disrupting inner hair cell-mediated activity in spiral ganglion cells alters their molecular specification ^89,90^. In the visual system, light-evoked activity controls vascular patterning in the retina ^91^, transcriptional maturation of retinal ganglion cells ^92^, and refinement of retinal connectivity ^93-95^. The role of vision in the maturation of retinal ganglion cell dendritic arbors is reminiscent of the relationship we report here between mechanically evoked activity and mechanosensory end organ structures.

Different stages of spontaneous and stimulus-evoked neuronal activity patterns are detected in DRG neurons during fetal and early postnatal development ^96,97^, though their contributions to somatosensory neuron development remain unknown. It is tempting to speculate that different temporally defined phases of activity could play distinct roles in the maturation of the somatosensory system, analogous to the waves of activity that pattern other sensory systems. Moreover, as work in adult mouse olfactory sensory neurons and *C. elegans* temperature sensitive neurons has demonstrated that activity-dependent gene expression can tune peripheral sensory neuron responses to changing environmental cues ^98-100^, it will be of interest to explore how different patterns of neuronal activity in primary mechanosensory neurons shape the maturation of their functional response properties during development and in mature circuits.

## Supporting information

Supplemental Figures 1-7 with legends, Supplemental Table 1

## Acknowledgements

We thank Erica Huey, Hankyul Kwak, Karina Lezgiyeva, Rosa Martinez-Garcia, Kitwa Ng, Lijun Qi, Zoe Sarafis, and Andrew Shuster for comments on the manuscript. We thank Tracy Tran, Carol Eisenberg, and Jack Delucia for help with preliminary analyses of Neuropilin-2 gene knock out animals. We thank Miriam H. Meisler for Scn8aflox mice and William A. Catterall for Scn1aflox mice. We thank the Neurobiology Imaging Facility at Harvard Medical School for assistance with tissue processing. This work was supported by NIH grants K99 NS124993 (CS), 1DP2NS127278 (NS), 1R01 AT011447-01 (DDG), R35 5R35NS097344-05 (DDG), a William Randolph Hearst Fellowship (CS), The Klingenstein-Simons Foundation, (NS), The Whitehall Foundation (NS), and the Edward R. and Anne G. Lefler Center for Neurodegenerative Disorders (DDG). DDG is an investigator of the Howard Hughes Medical Institute. This article is subject to HHMI’s Open Access to Publications policy. HHMI lab heads have previously granted a nonexclusive CC BY 4.0 license to the public and a sublicensable license to HHMI in their research articles. Pursuant to those licenses, the author-accepted manuscript of this article can be made freely available undera CC BY 4.0 license immediately upon publication.

## Author Contributions

CS and DDG conceived the study. CS, NA, JS, AH, AT, SR, and ARM performed anatomy experiments. NS performed RNA sequencing experiments. NS and CS analyzed RNA sequencing data. CS performed electrophysiology experiments with assistance from JT. RA assisted with experimental design and tissue collection. AH, MNT, and MI performed electron microscopy experiments. BPL and SP provided intellectual guidance on the project. CS and DDG wrote the manuscript with input from all authors.

## Methods

### Lead contact

Further information and requests for resources and reagents should be directed to and will be fulfilled by the lead contact, David Ginty

### Materials availability

This study did not generate new unique reagents.

### Data and code availability

All data reported in this study and all code will be shared by the lead contact upon request.

### Experimental Model Details

Mice used in the study were of mixed background (CD1 and C57BL/6J) except for *Rem2^Null^* mice, which were pure 129S2/SvPasCrl (Charles River). Previously published mice used in this study include: *Cdx2-Cre* (JAX# 009350)^41^, *Piezo2^null^*^25^, *Piezo2^fl^*^ox^ (JAX # 027720) ^19^, *Rosa26^LSL-tdTomato^* (Ai14)(JAX# 007914) ^101^, *Rosa26^LSL-synaptophysin-tdTomato^* (Ai34)(JAX# 012570)^102^, *Dhh^Cre^* (JAX# 012929)^46^, *Rem^Null^*^65^, *Avil^Cre^* ^103^, *Scn8a^flox^ ^104^*, *Scn1a^flox^* ^105^, *Ntrk2^tauEGFP^*^12^, *Na_v_1.8Cre* (JAX # 036564) ^51^. Mice were handled and housed in standard cages in accordance with the Harvard Medical School and IACUC guidelines. Both male and female mice were used in all experiments.

Control littermates of *Piezo2*, *Scn8a*, and *Scn1a* conditional mutants included animals of several different genotypes and are listed for each experiment in Supplemental Table 1. Our data indicated no difference among controls of different genotypes; therefore, they were combined for statistical analyses and collectively referred to as “Controls.” Likewise, *Cdx2-Cre/+; Piezo2^flox/Null^* and *Cdx2-Cre/+; Piezo2^flox/flox^* animals were indistinguishable in the severity of their morphological, behavioral, and electrophysiological phenotypes (see also ^32^) and were combined for statistical analyses.

### Method Details

#### Antibody staining of whole-mount tissue or cryosections

The following dilutions and primary antibodies and lectins were prepared: chicken anti-NFH (Aves # NFH, 1:500), rabbit anti-NF200 (Sigma # N4142, 1:500), rabbit anti-Tubulin beta 3 (TUBB3) (Biolegend #Poly18020, 1:500), rabbit anti-S100 Beta (VWR/Protein Tech #15146-1-AP, 1:300), mouse anti-NeuN (Sigma # MAB377, 1:500), rat anti-Troma (DSHB TROMA-I supernatant, 1:100), goat anti-GFP (US Biological Life Sciences # G8965-01E, 1:500), goat anti-mCherry (CedarLane # AB0040-200, 1:500), guinea pig anti-vGluT1 (Millipore #AB5905, 1:1000), rabbit anti-CGRP (Immunostar #24112, 1:500), mouse anti-Kv4.3 (Antibodies Inc. #75-017, 1:250), Isolectin B4 (Alexa 647 conjugated), Isolectin B4 (Alexa 488 conjugated). The secondary antibodies used were Alexa 488, 546 or 647 conjugated donkey or goat anti-mouse, rabbit, chicken, goat or guinea pig (Life Technologies or Jackson ImmunoResearch) and were prepared at 1:500 or 1:1000 dilutions.

For whole-mount staining of trunk hairy skin and Pacinian corpuscles, animals P5 and older were anesthetized under isoflurane and euthanized by cervical dislocation. Pups younger than P5 were anesthetized with ice and then decapitated. Hairy skin was treated with commercial depilatory cream (NAIR, Church and Dwight Co.) and then finely dissected, using a spatula to remove fat tissue underneath the skin. Skin was rinsed with 1xPBS and fixed for 1.5-2 hours in 2% paraformaldehyde in PBS or Zamboni’s fixative (Fisher NC9335034) at 4°C, then washed 3×10 minutes in PBS. Hairy skin was cut into small pieces (1 cm x1 cm) before antibody staining. For Pacinian corpuscle dissections, the tibia and fibula were dissected from the hind limb immediately after euthanasia, keeping muscle and surrounding tissue intact, and fixed overnight or 2 hours in 2% paraformaldehyde or Zamboni’s fixative at 4°C. Legs were washed 3×10 minutes in PBS, then finely dissected to remove the interosseous membrane from the bone. Interosseous membranes and hairy skin were then processed for antibody staining as described: Tissue was washed every 30 minutes for 5-6 hours with PBST (PBS + triton X-100). For hairy skin, 0.3% Triton X-100 in PBS was used. For Pacinians, 1% Triton X-100 in PBS was used. Primary antibodies were applied in blocking solution (5% normal goat or donkey serum, 75% PBST, 20% DMSO, 0.01% NaN_3_) and incubated at room temperature for 3-5 nights, with gentle rocking. Tissues were rinsed 3x in PBST, then washed every 30 minutes for 5-6 hours in PBST. Secondary antibodies were applied in blocking solution and incubated at room temperature for 2-3 nights, with gentle rocking. Tissues were rinsed 3x in PBST, washed every 30 minutes for 5-6 hours in PBST, then serially dehydrated (10-15 minutes per wash) in 50%, 75%, and 100% ethanol diluted in distilled water. Tissue was stored at -20°C in 100% ethanol for up to 5 days before clearing and imaging. Tissue was cleared in BABB (1 part benzyl alcohol to 2 parts benzyl benzoate) for 1-2 hours at room temperature and mounted in BABB for imaging. For mounting, chambers were made on slides consisting of a thin layer of vacuum grease to create four walls followed by placement of a coverslip. Tissue was imaged on a Zeiss LSM 700 confocal microscope using 10x or 20x objectives.

For immunohistochemistry of skin, DRG, and spinal cord cryosections, hairy or glabrous skin was finely dissected from animals 1-2 minutes after euthanasia or transcardial perfusion with 4% paraformaldehyde in PBS. Hair was removed with a commercial depilatory cream (Nair, Church & Dwight). Tissue was fixed (or post-fixed if mice were perfused first) 1.5-2 hours in 2% paraformaldehyde or Zamboni’s fixative at 4°C with gentle rotation, then washed 3×10 minutes in PBS. For spinal cord/DRG collection, animals were transcardially perfused with 4% paraformaldehyde in PBS. The spinal column was dissected out and drop-fixed in 4% PFA in PBS overnight. Tissue was washed 3×10 minutes in PBS, and spinal cords and DRG were finely dissected. Tissues were cryoprotected in 30% sucrose in 1x PBS at 4°C for 1-2 days, embedded in OCT (Fisher 1437365), frozen on a bed of dry ice, and stored at -80°C. Tissues were cryosectioned (25μm) using a cryostat (Leica) and collected on glass slides (12-550-15, Fisher). Sections were washed 3×5 minutes with 1xPBS containing 0.1% Triton X-100 (0.1% PBST), incubated with blocking solution (0.1% PBST containing 5% normal goat serum (S-1000, Vector Labs) or normal donkey serum (005-000-121, Jackson)) for 1 hour at RT, incubated with primary antibodies diluted in blocking solution at 4°C for 1-2 nights, washed 3×10 minutes with 0.1% PBST, incubated with secondary antibodies diluted in blocking solution 2 hours at room temperature, washed again 4×10 minutes with 0.1% PBST, and mounted in Fluoromount-G mounting medium (Southern Biotech 0100-01) or DAPI-Fluoromount-G (Southern Biotech 0100-20).

Two antibodies against neurofilament heavy chain protein were used in this study: Chick anti-NFH (Aves #NFH) was used for whole-mount staining of Pacinian corpuscles and for staining Meissner corpuscles, and rabbit anti-NF200 (Sigma # N4142) was used for whole-mount staining of trunk hairy skin.

#### Perfusions

Mice were anesthetized under isoflurane and transcardially perfused with 5-10 mL of Ames Media (Sigma) in PBS with heparin (10 U/mL), followed by 5-10 mL of 4% paraformaldehyde (PFA) in PBS. After perfusion, the skull, vertebral column, and central nervous system were removed and post-fixed in 4% PFA in PBS at 4°C overnight. Samples were washed 3×10 min in PBS at room temp after overnight fixation before fine dissection.

#### Skin Injections

Mice P3-P4 were anesthetized using ice for 2 to 3 minutes. Mice P7 and older were anesthetized with continuous inhalation of 2% isoflurane from a precision vaporizer for the duration of the procedure (5-10 min). The animal’s breathing rate was monitored throughout the procedure and the anesthetic dose was adjusted as needed. The skin region of interest (dorsal or ventral hind paw) was swabbed with ethanol. Injections were done with a beveled borosilicate pipette. Forceps were used to stabilize the skin while the needle was inserted into the dermis, injecting 0.5-2 μL of virus. Mice were injected in 1-6 locations within the skin region of interest, depending on skin identity and the amount of labeling desired. For AAV-Cre-mediated deletion of *Piezo2*, mice harboring a Cre dependent reporter allele and *Piezo2^flox/Null^* alleles were injected with AAV2-retro-hSyn-Cre (1.2e13, Addgene 105553) at P3-P4 on both sides of the paw. For sparse labeling of Aβ-LTMR axon terminals in the dorsal column nuclei, wild type or *Cdx2-Cre; Piezo2 cKO* mice age P14 or older were injected with AAV2/retro-synaptophysin-TdTomato (1e14, UPenn Viral Core) or AAV2/retro-hSyn-eGFP (1.7e13, Addgene 50465), to either the dorsal hairy or ventral glabrous paw. For all injections, a small amount of fast green (Sigma F7252-5G) in 0.9% saline was added to the virus. After injection, mice recovered from anesthesia on a warm pad while being monitored. Mice were returned to the cage once normal activity had resumed. Five weeks post-injection, animals were sacrificed by transcardial perfusion under isoflurane anesthesia. For sparse labeling of Aβ-LTMRs with syn-tdTomato or GFP, animals were screened following tissue collection to confirm that fluorescent protein expression was restricted to DRGs at somatotopically appropriate segmental levels (Lumbar level L2-L5, unilateral signal on the injected side).

#### Electron microscopy imaging of Meissner corpuscles

Forepaw or hindpaw digit tips were isolated and immersed in a glutaraldehyde/formaldehyde fixative for 1 hour at room temperature, further dissected to remove muscle and fat, and subsequently fixed overnight at 4°C. Sample preparation was done as previously described ^106^. Ultrathin sections were cut at 50-70 nm and imaged using a JEOL 1200EX transmission electron microscope at 80 kV accelerating voltage. Images were cropped and adjusted to enhance contrast using Fiji/ImageJ, and pseudocolored by hand.

#### End-organ analysis

For Meissner corpuscle (MC) analysis, hind paw plantar pads were sectioned, stained, and imaged as described above. Images were analyzed using FIJI. Maximum projections of z stacks were generated. DAPI was used to identify the dermal papillae structures, inside which MC are located. Neurofilament (NFH) and S100 were used to identify the MC. The selection tool on FIJI was used to draw the outline of the S100^+^ corpuscle located inside the dermal papillae and the Measure tool was used to calculate its area and width. To quantify NFH^+^ innervation of corpuscles, the number of NFH^+^ axons innervating MCs in the image was divided by the number of MCs in the image. To quantify NFH^+^ or S100^+^ protrusions from the MC into the epidermis, the number of NFH^+^ or S100^+^ protrusions from all MCs in the image was divided by the number of MCs in the image.

For Pacinian corpuscle (PC) analysis, PCs from the interosseous membrane surrounding the fibula were dissected, stained, and imaged as described above. Images were analyzed using FIJI. For each animal, the total number of PCs identified in the interosseous membrane dissection was quantified. Each PC was scored as belonging to one of three categories: 1) innervated by an unbranched NFH^+^ axon, 2) innervated by an NFH^+^ axon that branches outside of the ultraterminal region, 3) innervated by no NFH^+^ axon. To compare the percentage of PCs containing branched axons, animals were only included in statistical analyses if at least eight PCs were detected after interosseous membrane dissection and staining. To measure Pacinian corpuscle areas and lengths, maximum projections of Pacinian corpuscles were generated on FIJI, and two measurements were obtained: First, the S100^+^ area was traced using the polygon tool. Second the length of the S100^+^ area of the Pacinian corpuscle (in the same direction as the length of the NFH+ axon) was measured using the line tool.

For hairy skin peripheral endings, trunk hairy skin was collected, stained, and imaged as described above. Images were analyzed using FIJI. To quantify the percentage of hair follicles innervated by TUJ1^+^ or NFH/NF200^+^ lanceolate endings, maximum projections were generated and the number of hair follicles in each image was quantified. 3-4 images were analyzed per animal, and the average value was recorded for that animal. To quantify the number of NFH/NF200^+^ lanceolate endings around each hair follicle, Troma was used to identify guard from non-guard hair follicles, and individual NFH/NF200^+^ lanceolate endings were counted using the Cell Counter plugin on FIJI. For a subset of animals, NFH/NF200^+^ lanceolate endings were reconstructed using the SNT plugin, and maximum projections of filled neurons were generated. To quantify Merkel cell innervating LTMRs at touch domes, guard hair follicles were identified using Troma staining, and the number of Troma^+^ cells per touch dome was quantified using the Cell Counter plugin. The number of NFH/NF200^+^ axon-Merkel cell contacts was also quantified for each touch dome. To quantify the percentage of hair follicles innervated by CGRP^+^ or NFH^+^ lanceolate endings, maximum projections were generated and the number of hair follicles in each image was quantified. 3-4 images were analyzed per animal, and the average value was recorded for each animal. For all conditions, tissue was processed and imaged in parallel, with at least one control and one mutant processed together using the same reagents. Images were analyzed blind to genotype.

#### Spinal cord analysis

Spinal cords were sectioned, stained, and imaged as described above. Images were analyzed using FIJI. Maximum projections of z stacks were generated, and the height of the spinal cord dorsal horn was measured. Then, the height of the analyzed region of interest (CGRP^+^, vGlut1^-^, or IB4^+^) was measured. Images were analyzed blind to genotype; for each animal, 4 or more images were analyzed, and the average values were recorded.

#### DRG marker analysis

DRG were sectioned, stained, and imaged as described above. Images were analyzed using FIJI. Maximum projections of z stacks were generated, and the number of cells positive for each marker (IB4, CGRP, NF200, Th, *TrkB^GFP^,* K_v_4.3) was quantified using the Cell Counter plugin. Images were analyzed blind to genotype; for each animal, 3 or more images were analyzed, and the average values were recorded.

#### Immunohistochemistry of free-floating brainstem sections

The brain, brainstem, and spinal cord were removed from the skull and vertebral column. The skull was cut down the midline and pulled laterally to remove. The dorsal and ventral sides of the vertebral column were removed. Then, forceps were used to remove the dorsal root ganglia (DRG) bilaterally from the vertebral column and surrounding tissue, retaining the connection to the spinal cord. This was done for all DRG working from lumbar to cervical levels. Once all DRG had been removed, the brain, brainstem, and spinal cord could be extracted from the vertebral column and stored in PBS. Sectioning of tissue samples was done using a vibrating blade microtome (Leica VT100S). Samples were embedded in 3% agarose in 1x PBS. Once solidified, samples were trimmed and attached to the sectioning plate with superglue. Transverse brainstem and spinal cord sections were taken at 100 μm while horizontal brain sections were taken at 200 μm. Tissue samples were then rinsed in 50% ethanol/water solution for 30 minutes, followed by 3×10 min in 2xPBS. Tissue samples were incubated in a mixture of primary antibodies in high salt Phosphate Buffer Saline (2x PBS) containing 0.1% Triton X-1000 (2xPBST) for 72 hours at 4°C. Tissue was rinsed 3×10 min with 2xPBS before incubation in the secondary antibody solution for 24 hours at 4°C. The secondary antibody solution was made up in 2xPBST and contained species-specific Alexa Fluor 488, 546, and 647 conjugated IgGs. The tissue was then mounted on glass slides, coverslipped in VectaShield mounting medium (Vector labs H-1000) and stored at 4°C until imaging.

#### Analysis of pre-synaptic boutons in the GN

To label pre-synaptic boutons of cutaneous sensory neurons, *R26^syt-tdTomato^* (Ai34) animals were injected with AAV2/retro-Cre virus, and wild type animals (CD1) or *Cdx2-Cre; Piezo2 cKO* animals were injected with either AAV2-retro-synaptophysin-tdTomato or AAV2-retro-hSyn-eGFP into the hairy or glabrous paw dermis at age P14 or older. Animals were transcardially perfused with 4% paraformaldehyde 5-6 weeks post injection. After overnight post-fixation in 4% PFA at 4°C and three washes in 1xPBS, dorsal column nuclei were finely dissected and sectioned transversely on the vibratome at 100 µm. Antibody staining was performed as described above. Sections were mounted in VectaShield and imaged on a Zeiss LSM 700 confocal microscope using a 40x oil-immersion lens. FP^+^ puncta in the DCN were identified manually using ImageJ. Automatic thresholding and background subtraction were applied to all images. Co-localization with vGlut1 puncta was assessed manually after bouton identification.

#### DRG dissection for RNA sequencing

DRG dissection was performed as described ^54^, except that ganglia above cervical level 2 were excluded. Briefly, animals were sacrificed and spinal columns rapidly removed and placed on ice. Individual DRG with central and peripheral nerves attached were removed from axial levels below C2 and placed into ice-cold DMEM:F12 (1:1) (Gibco 11330-032) supplemented with 1% pen/strep and 12.5 mM D-glucose. A fine dissection was performed to remove the peripheral and central nerve roots, resulting in only the sensory ganglia remaining. scRNA-seq experiments are the culmination of two independent bioreplicates. In the first bioreplicate, DRG from two *Cdx2-Cre/+; Piezo2^flox/flox^* animals (age P24) were dissected and assigned as “Mutant sample 1” and DRG from three control littermates were dissected on the same day and assigned as “Control sample 1.” In the second bioreplicate, DRG from three *Cdx2-Cre/+; Piezo2^flox/flox^* animals (age P21) were dissected and assigned as “Mutant sample 2” and DRG from two control littermates were dissected and assigned as “Control sample 2.” Sensory ganglia were dissociated in 40 units papain, 4 mg/ml collagenase, 10 mg/ml BSA, 1 mg/ml hyalurdonidase, 0.6 mg/ml DNase in DMEM:F12 + 1% pen/strep + 12.5 mM glucose for 25 min at 37 °C. Digestion was quenched using 20 mg/ml ovomucoid (trypsin inhibitor), 20 mg/ml BSA in DMEM:F12 + 1% pen/strep + 12.5 mM glucose. Ganglia were gently triturated with fire-polished glass pipettes (opening diameter of approx. 150–200 μm). Neurons were then passed through a 70-μm filter to remove cell doublets and debris. Neurons were pelleted and washed 4 times in 20 mg/ml ovomucoid (trypsin inhibitor), 20 mg/ml BSA in DMEM:F12 + 1% pen/strep + 12.5 mM glucose followed by 2× washes with DMEM:F12 + 1% pen/strep + 12.5 mM glucose all at 4 °C. After washing, cells were resuspended in 45 μl of DMEM:F12 + 1% pen/strep + 12.5 mM glucose.

#### Single cell RNA sequencing analysis

Alignment, mapping, and quality control was performed using the 10x genomics cell ranger pipeline. This pipeline generated the gene expression tables for individual cells used in this study. Briefly, ∼8000-10000 dissociated cells from DRG were loaded per 10x run (10x genomics chromium single cell kit, v3). Downstream reverse transcription, cDNA synthesis and library preparation were performed according to manufacturer’s instructions. All samples were sequenced on a NextSeq 550 with 58bp sequenced into the 3’ end of the mRNAs. For quality control filtering, individual cells were removed from the dataset if they had fewer than 1,000 discovered genes, fewer than 1,000 unique molecule identifiers (UMIs) or more than 12% of reads mapping to mitochondrial genes. Non-neuronal cells were also removed from our analysis: first, by identifying clusters with prominent expression of non-neuronal markers (*Glul*, *Plp1*, *Dhh*, *Fcgr3*) and removing them; second, by checking for DRG sensory neuron markers (*Pou4f1*, *Avil*, *Tubb3*) and removing any clusters that did not express them. Neuronal subtypes were identified as clusters using PCA/UMAP analysis combined with prior studies identifying marker genes for distinct DRG neuron subtypes. Control and *Cdx2 Piezo2 cKO* cells from each bioreplicate were merged for downstream analysis. Differential gene expression analysis was performed on all expressed genes using the FindMarker function in Seurat using the Wilcoxon Rank Sum test for a log fold threshold change of 0.5 or greater. A pseudocount of 0.001 was added to each gene to prevent infinite values. Adjusted p values <0.05 were considered significant.

#### *In situ* hybridization

Detection of transcripts was performed by fluorescent i*n situ* hybridization. Individual DRG ganglia were rapidly dissected from euthanized mice and frozen in dry ice cooled 2 methylbutane, then stored at -80°C until further processing. DRGs were cryosectioned at a thickness of 20 μm and RNA was detected using RNAscope (Advanced Cell Diagnostics) according to the manufacturer’s protocol. The following probes were used: Mm Piezo2 exons 43-45 (Cat# 439971-C3), Mm-Nefh (Cat# 443671-C2), tdTomato (Cat# 317041), Cre (Cat#533031), Mm-Th (Cat#317621-C2), Mm-Rem2 (Cat#501801), Mm-Nrp2 (Cat#500661), Mm-Colq (Cat# 496211-C2), Mm-Calb1 (Cat# 428431-C3). Sections were mounted in FluoroMount-G (Fisher 0100-01) and imaged on a Zeiss LSM 700 confocal microscope using a 20x air objective.

#### Analysis of gene expression after *in situ* hybridization

Tissue was stained and imaged as described above. For quantification of Piezo2^+^/Nefh^+^ cells, images were analyzed on FIJI. Maximum projections of z stacks were generated, and Nefh^+^, tdTomato^+^, Cre^+^, and Piezo2^+^ cells were counted using the Cell Counter plugin. For each animal, 5 or more images were analyzed, and the average values were recorded. For quantification of Rem2^+^ cells, images were analyzed on FIJI. Maximum projections of z stacks were generated, and Calb1^+^ cells were counted using the Cell Counter plugin. Then, the number of Calb1^+^/Rem2^+^ cells were counted. For each animal, 5 or more images were analyzed, and the average values were recorded.

#### *In vivo* electrophysiological recordings

Juxtacellular recordings were done on DRG neurons *in vivo*. All experiments were performed blind to genotype. Mice of both sexes age P21-P30 were anesthetized with urethane (1.5 g/kg). In some cases, mice encountered difficulty breathing following urethane administration, and a tracheotomy was performed to improve survival rates. The skin and muscle above the trachea were dissected and a small incision was made in the trachea; a plastic tube was fixed in place with superglue to allow breathing to continue. To expose DRG, an incision was made over the lumbar spine. The L4 DRG was exposed using fine dissection forceps, and two custom made spinal clamps fixed the spine in place. A glass electrode (borosilicate, 1-2 MO) filled with saline was lowered into the DRG. To search for mechanically sensitive units, a custom-built mechanical stimulator delivered a 200 Hz or 300 Hz vibration stimulus (∼20 mN) to the middle of the leg, above the tibia. Receptive fields of units were then identified through an iterative process in which the stimulator was moved to different locations while reducing the force of the vibration stimulus. Gentle brushing with a paintbrush or cotton swab was also used to identify receptive fields. Units were then characterized based on their responses to air puff, indentation, and vibration (10-500 Hz, 0-35 mN). Signals were amplified using a Multiclamp 700 A/B and acquired using a Digidata 1550A/B using Clampex 10/11 software.

#### DRG electrophysiology analysis

Units were sorted into categories based on their response properties. Units that had their receptive field on hairy skin and were activated by movement of hairs using fine forceps or air puff were categorized as “hairy.” Units that did not have a well-defined cutaneous receptive field but could become phase-locked to high frequency vibration (200 Hz or higher) at forces ≤ 10 mN were classified as “Pacinian”. The best receptive fields of these units were often on the heel or ankle region, though they were very large and could be activated by mechanical stimulation delivered to most areas of the hindlimb. Units that did not have a cutaneous receptive field but responded strongly to moving the leg or paw were classified as “Proprioceptors.” Proprioceptors units from all genotypes often had high levels of background firing. No units with receptive fields on glabrous paw skin were identified using this search stimulus. Data were analyzed on Clampfit and Matlab. Indentation threshold was defined as the minimum force required for a unit to be activated by an indentation stimulus, across several sweeps. Each force was tested 4-6 times per unit. Maximum firing rates for hairy skin LTMRs were obtained during 20 second mechanical stimulus intervals during which the skin was lightly brushed or received an air puff stimulus delivered from a canister. Both air puff and brush were tested for all hairy skin LTMRs, and the minimum interstimulus interval (ISI) between 3 consecutive spikes was recorded for each trace. Maximum firing rates were calculated, and the highest value obtained for each neuron in response to any mechanical stimulus (either brush or air puff) was recorded. For analysis of responses to vibration, threshold for entrainment was defined as the minimum force required for a unit to reach 50% entrainment (i.e., firing every other cycle or more) or 100% entrainment (i.e., firing every cycle or more). Forces were tested 3-8 times for each unit, and the minimum force able to elicit entrainment was recorded.

#### Statistical Analysis

Statistical tests were conducted using GraphPad Prism or R. Normality of the data was tested using the Shapiro-Wilk test and equal variances were tested using the F test. Comparisons between two independent groups were performed using the unpaired t test (in the case of two groups distributed normally, with no significant difference between variances), Welch’s t test (in the case of two groups distributed normally, but with unequal variances), or Mann-Whitney test (non-parametric data). Comparisons among more than 2 independent groups were performed using one-way ANOVA (parametric distribution for all groups) or the Kruskal-Wallis test (non-parametric distribution for at least one group), and multiple comparisons were performed using the post-hoc test indicated in the figure legend. An adjusted p value < 0.05 was considered significant.

