## Supplemental Figures 1-7 with legends, Supplemental Table 1 for "Activity-dependent development of the body’s touch receptors"

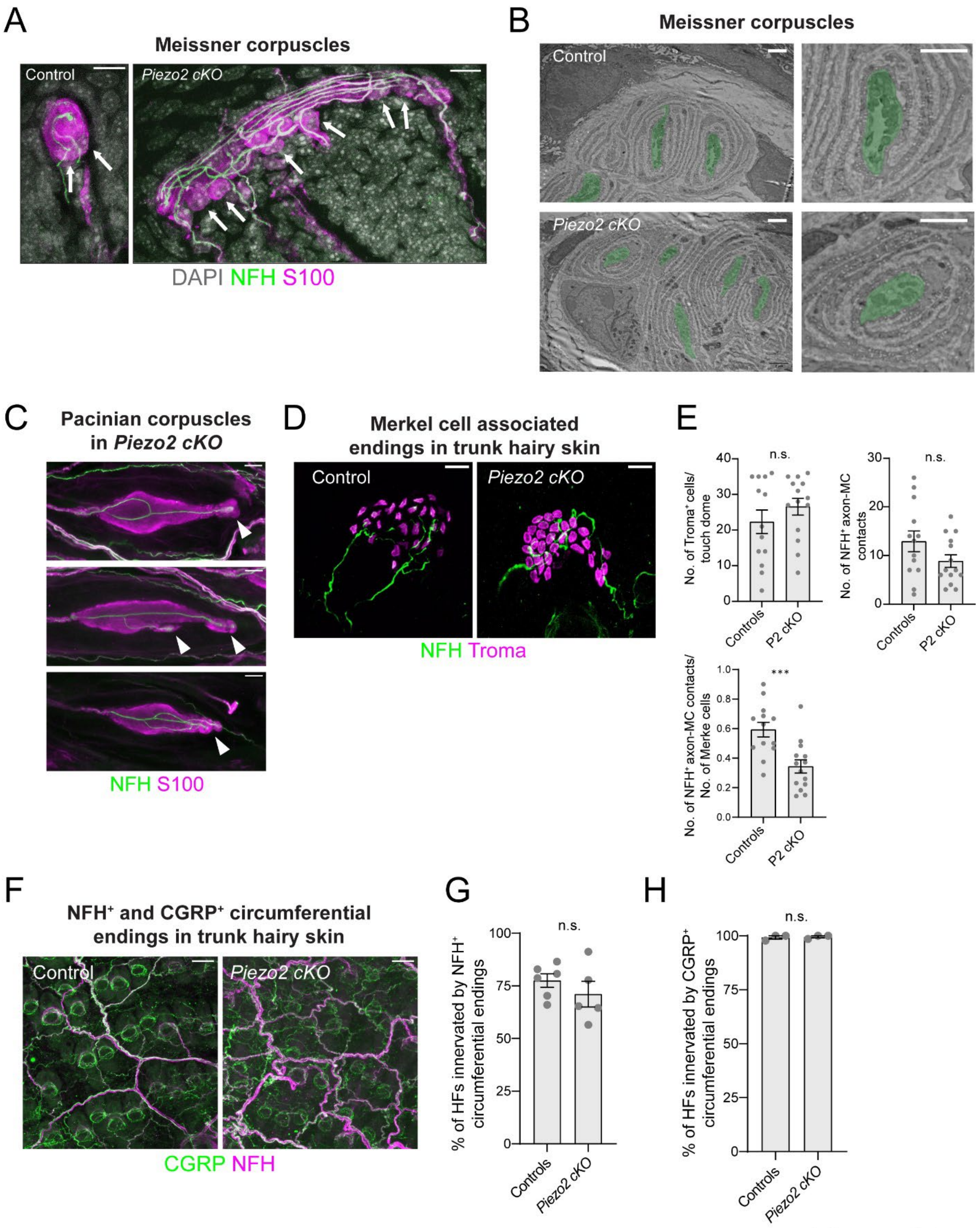

Figure S1

**Supplemental Figure 1: Additional characterization of LTMR structures in *Cdx2-Cre; Piezo2 cKO* mutants, related to Figure 1**

**A.** Left: High magnification confocal microscopy images of Meissner corpuscles in the hind paw plantar pads of a control littermate (+/+; *Piezo2*$^{+/+}$) or a *Cdx2-Cre; Piezo2*$^{flox/Null}$ mutant. Arrows indicate S100$^{+}$/DAPI$^{+}$ nuclei of lamellar cells within the corpuscles. Scale bar = 10 µm.

**B.** Electron microscopy images of sections through Meissner corpuscles in the hind paw finger pads of a control littermate or a *Cdx2-Cre/+; Piezo2*$^{flox/Null}$ mutant, showing normal ultrastructural organization. Axon profiles are outlined and colored in green; insets on the right show two axons. The experiment was repeated in hind paw tissue collected from one additional *Cdx2-Cre/+; Piezo2*$^{flox/Null}$ animal, and in forepaw tissue from one *Cdx2-Cre; Piezo2*$^{flox/flox}$ animal, with similar results. Scale bar = 2 µm.

**C.** Additional examples of Pacinian corpuscles from the hind limbs of a *Cdx2-Cre/+; Piezo2*$^{flox/flox}$ mutant. Scale bar = 20 µm. Arrowheads indicate the ultraterminal regions.

**D.** Images of touch domes found at guard hair follicles in trunk hairy skin of a control littermate or *Cdx2-Cre; Piezo2 cKO* animal. In animals from both groups, NF200$^{+}$ axons (green) form small extensions that interact with Troma$^{+}$ Merkel cells (magenta). Scale bar = 20 µm.

**E.** Quantification of touch dome morphology. Top, left: Number of Troma$^{+}$ Merkel cells/ touch dome. There was no difference between controls and *Cdx2-Cre; Piezo2 cKO* mutants (unpaired t-test). Top, right: Number of NF200$^{+}$ axon-Merkel cell contacts per touch dome. There was no difference between controls and *Cdx2-Cre; Piezo2 cKO* mutants (unpaired t-test). Bottom: Number of Nf200$^{+}$ axon-Merkel cell contacts divided by the total number of Merkel cells at a touch dome. There was a reduction in *Cdx2-Cre; Piezo2 cKO* mutants (***$p<0.001$, unpaired t-test). For all three plots, each data point indicates one touch dome at a guard hair follicle; averages are shown; error bars indicate the s.em. 3 controls and 3 *Cdx2-Cre; Piezo2 cKO* mutants were analyzed.

**F.** Representative images of trunk hairy skin from a control animal or *Cdx2-Cre; Piezo2 cKO* animal, stained with anti-CGRP (green) or anti-NFH (purple) antibodies. Scale bar = 50 µm.

**G.** The percentage of all hair follicles (HFs) innervated by NFH$^{+}$ circumferential endings in trunk hairy skin is shown. There was no difference between controls and *Cdx2-Cre; Piezo2 cKO* animals (unpaired t test). Each data point indicates one animal; averages are plotted; error bars indicate s.e.m.

**H.** The percentage of all hair follicles (HFs) innervated by CGRP$^{+}$ circumferential endings in trunk hairy skin is plotted. There was no difference between controls and *Cdx2-Cre; Piezo2 cKO* animals (Mann-Whitney U test). Each data point indicates one animal; averages are plotted; error bars indicate s.e.m.

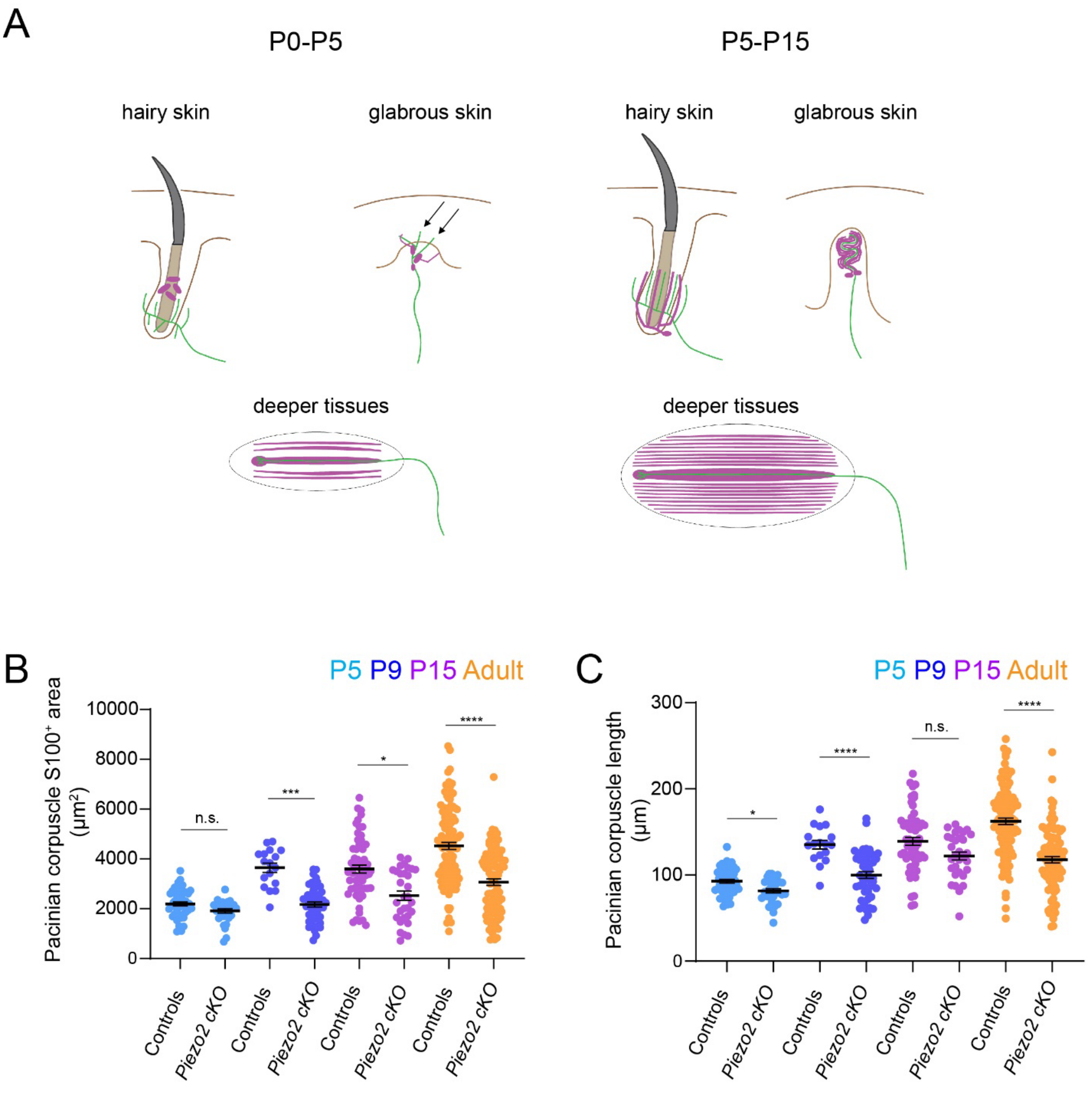

Figure S2

**Supplemental Figure 2: Additional characterization of mechanosensory end organs over development, related to Figure 2**

**A.** Summary of mechanosensory end organ development in rodents. In the first few days after birth, Aβ sensory fibers (green) are found associated with guard hair follicles in hairy skin and have innervated the dermal papillae of glabrous skin, where some nerve fibers and their accompanying Schwann cells (purple) extend immature processes that invade the epidermis (arrows). At this stage, immature Pacinian corpuscles can be found in the interosseous membrane surrounding bones: they consist of a single unbranched axon (green) enclosed in loosely arranged lamellar cells (purple) that form the immature inner and outer cores of the corpuscle. Over the first two postnatal weeks, mechanosensory structures associated with Aβ axons mature until they reach their adult-like morphologies at around P15.

**B.** Quantification of Pacinian corpuscle areas over development. The $S100^+$ areas of Pacinian corpuscles were measured. There was no significant difference between control and *Cdx2-Cre; Piezo2 cKO* mutants at P5, but Pacinian corpuscles of *Cdx2-Cre; Piezo2 cKO* had smaller areas at P9, P15, and in adult animals (>P30) (*p<0.05, ***p<0.001, ****p<0.0001, Kruskal-Wallis test with post-hoc Dunn test). Each point indicates one corpuscle; averages are shown; error bars indicate the s.e.m. The following number of animals were analyzed: n=2 for P5 controls, n=2 for P5 *Piezo2 cKO*, n=2 for P9 controls, n=3 for P9 *Piezo2 cKO*, n=3 for P15 controls, n=2 for P15 *Piezo2 cKO*, n=8 for adult controls, n=8 for adult *Piezo2 cKO*. P5, P9, and P15 data are from the same animals analyzed in Figure 2A-B. Adult data include Pacinian corpuscles from the same animals analyzed in Figure 1B.

**C.** Quantification of Pacinian corpuscle lengths over development. The lengths of Pacinian corpuscles were measured. There was no significant difference between control and *Cdx2-Cre; Piezo2 cKO* mutants at P15, but Pacinian corpuscles of *Cdx2-Cre; Piezo2 cKO* had smaller lengths at P5, P9, and in adult animals (>P30) (*p<0.05, ****p<0.0001, Welch's ANOVA test with post-hoc Dunnett's T3 multiple comparisons test). Each point indicates one corpuscle; averages are plotted; error bars indicate the s.e.m. The same animals were analyzed as in Supplemental Figure 2B.

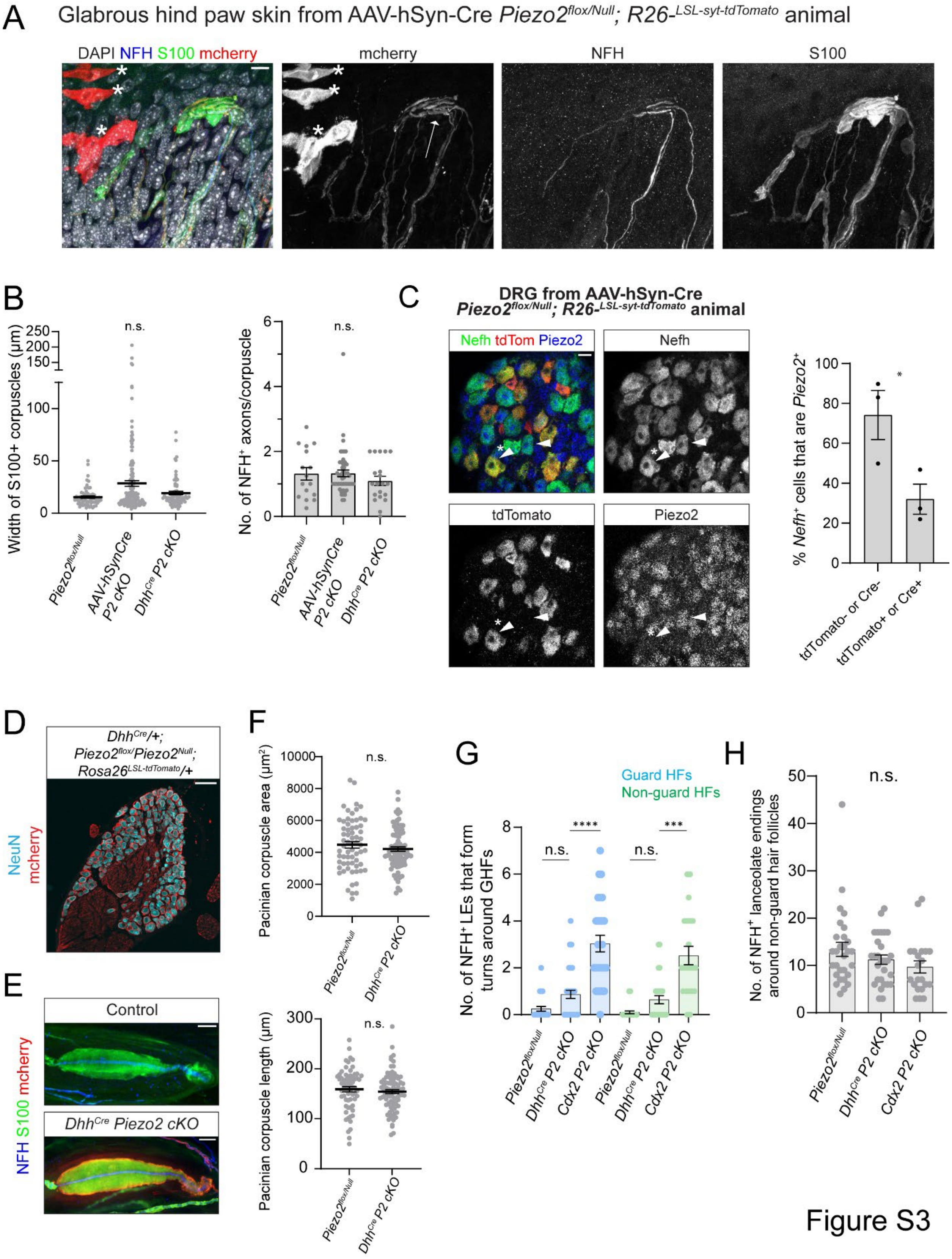
A
Glabrous hind paw skin from AAV-hSyn-Cre Piezo2flox/Null; R26-LSL-syt-tdTomato animal
DAPI NFH S100 mcherry
mcherry
NFH
S100
B
Width of S100+ corpuscles (µm)
n.s.
No. of NFH+ axons/corpuscle
Piezo2flox/Null
AAV-hSynCre P2 cKO
DhhCre P2 cKO
C
DRG from AAV-hSyn-Cre Piezo2flox/Null; R26-LSL-syt-tdTomato animal
Nefh tdTom Piezo2
Nefh
tdTomato
Piezo2
% Nefh+ cells that are Piezo2+
tdTomato- or Cre-
tdTomato+ or Cre+
D
DhhCre/+; Piezo2flox/Piezo2Null; Rosa26LSL-tdTomato/+
NeuN mcherry
E
Control
DhhCre Piezo2 cKO
NFH S100 mcherry
F
Pacinian corpuscle area (µm2)
Pacinian corpuscle length (µm)
DhhCre P2 cKO
G
Guard HFs
Non-guard HFs
No. of NFH+ LEs that form turns around GHFs
Cdx2 P2 cKO
H
No. of NFH+ lanceolate endings around non-guard hair follicles

Figure S3

**Supplemental Figure 3: Additional characterization of AAV-hSyn-Cre *Piezo2 cKO* and *Dhh$^{Cre}$ Piezo2 cKO* animals, related to Figure 3**

**A.** Images of a Meissner corpuscle from an AAV2/retro-hSyn-Cre -injected *Piezo2$^{flox/Null}$; Rosa26$^{LSLtdTomato/+}$* animal. Asterisks indicate labeling of non-neuronal cells in the epidermis. The arrow points to mcherry$^{+}$/NFH$^{+}$ axons innervating the corpuscle. Skin tissue was stained with antibodies against NFH (blue), S100 (green), and mcherry (red). DAPI staining is shown in gray. Scale bar = 10 µm.

**B.** Left: Quantification of the widths of S100$^{+}$ Meissner corpuscles in controls, AAV-hSyn-Cre injected *Piezo2$^{flox/Null}$* animals, and *Dhh$^{Cre}$/+; Piezo2$^{flox/Null}$* animals. There was no significant difference among the three groups (Kruskal-Wallis test with post-hoc Dunn test for multiple comparisons). Each point indicates one Meissner corpuscle; averages are shown by horizontal bars; error bars indicate the s.e.m. Data were collected from the same animals as in Figure 3D-E. Right: Quantification of the number of NFH$^{+}$ axons innervating each Meissner corpuscle divided by the total number of Meissner corpuscles in an image. There was no significant difference among the groups (Kruskal-Wallis test with post-hoc Dunn test). Each point indicates one image; averages are shown by horizontal bars; error bars indicate the s.e.m. Data were collected from the same animals as in Figure 3D-E.

**C.** Left: Lumbar-level DRG sections from an AAV2/retro-hSyn-Cre -injected *Piezo2$^{flox/Null}$; Rosa26$^{LSLtdTomato/+}$* animal, showing RNAscope signal for probes against *Nefh* (green), *tdTomato* (red), or *Piezo2* exons 43-45 (blue). Arrowhead indicates a *Piezo2$^{+}$/Nefh$^{+}$/tdTomato$^{-}$* cell. Arrowhead with asterisk indicates a *Piezo2$^{-}$/Nefh$^{+}$/tdTomato$^{+}$* cell. Scale bar = 25 µm. Right: % of *Nefh+* cells that are positive for *Piezo2*, grouped by whether the cells also express *tdTomato* or *Cre*. Each point indicates data from one animal; averages are shown; error bars indicate s.e.m. *$p<0.05$, unpaired t test.

**D.** Lumbar-level DRG section from *Dhh$^{Cre/+}$; Piezo2$^{flox/Null}$; Rosa26$^{LSLtdTomato/+}$* animal. Anti-mcherry signal (red) is detected in the satellite glial cells but is not present in the NeuN+ neurons (cyan). Scale bar = 100 µm.

**E.** Images of Pacinian corpuscles from +/+*; Piezo2$^{flox/Null}$* or *Dhh$^{Cre/+}$; Piezo2$^{flox/Null}$; Rosa26$^{LSL-tdTomato/+}$* animals. Tissue was stained with antibodies against NFH (blue), S100 (green), and mcherry (red). Scale bar = 20 µm.

**F**. Top: Quantification of Pacinian corpuscle areas in adult animals (>P30). There was no significant difference between +/+*; Piezo2$^{flox/Null}$* controls and *Dhh$^{Cre/+}$; Piezo2$^{flox/Null}$* mutants (Welch's t test). The following animals were analyzed: n=4 for controls, n=5 for *Dhh$^{Cre}$ Piezo2 cKO*. The data are from a subset of the animals analyzed in Figure 3F. Bottom: Quantification of Pacinian corpuscle lengths in adult animals (>P30). There was no significant difference between +/+*; Piezo2$^{flox/Null}$* controls and *Dhh$^{Cre/+}$; Piezo2$^{flox/Null}$* mutants (unpaired t test). The following animals were analyzed: n=4 for controls, n=5 for *Dhh$^{Cre}$ Piezo2 cKO*. The data are from a subset of the animals analyzed in Figure 3F.

**G.** Quantification of NFH$^{+}$ axonal structures formed by lanceolate endings around guard hair (blue) or non-guard hair follicles (green). Lanceolate endings in *Dhh$^{Cre/+}$; Piezo2$^{flox/Null}$* animals did not form more turns than those in *Piezo2$^{flox/Null}$* controls, and formed fewer turns than lanceolate endings in *Cdx2-Cre; Piezo2 cKO* animals (Kruskal-Wallis test with post-hoc Dunn multiple comparisons test, adjusted ****p<0.0001 and ***p=0.001). Control and *Dhh*$^{Cre}$ *cKO* data were collected from the same animals as in Figure 3H. Data for *Cdx2-Cre; Piezo2 cKO* animals are re-plotted from Figure 1F.

**H.** Quantification of the number of NFH$^{+}$ lanceolate endings around non-guard hair follicles. There was no significant difference among any of the groups (Kruskal-Wallis tested with post-hoc Dunn multiple comparisons tests). Control and *Dhh*$^{Cre}$ *cKO* data were collected from the same animals as in Figure 3H. Data for *Cdx2-Cre; Piezo2 cKO* animals are re-plotted from Figure 1F.

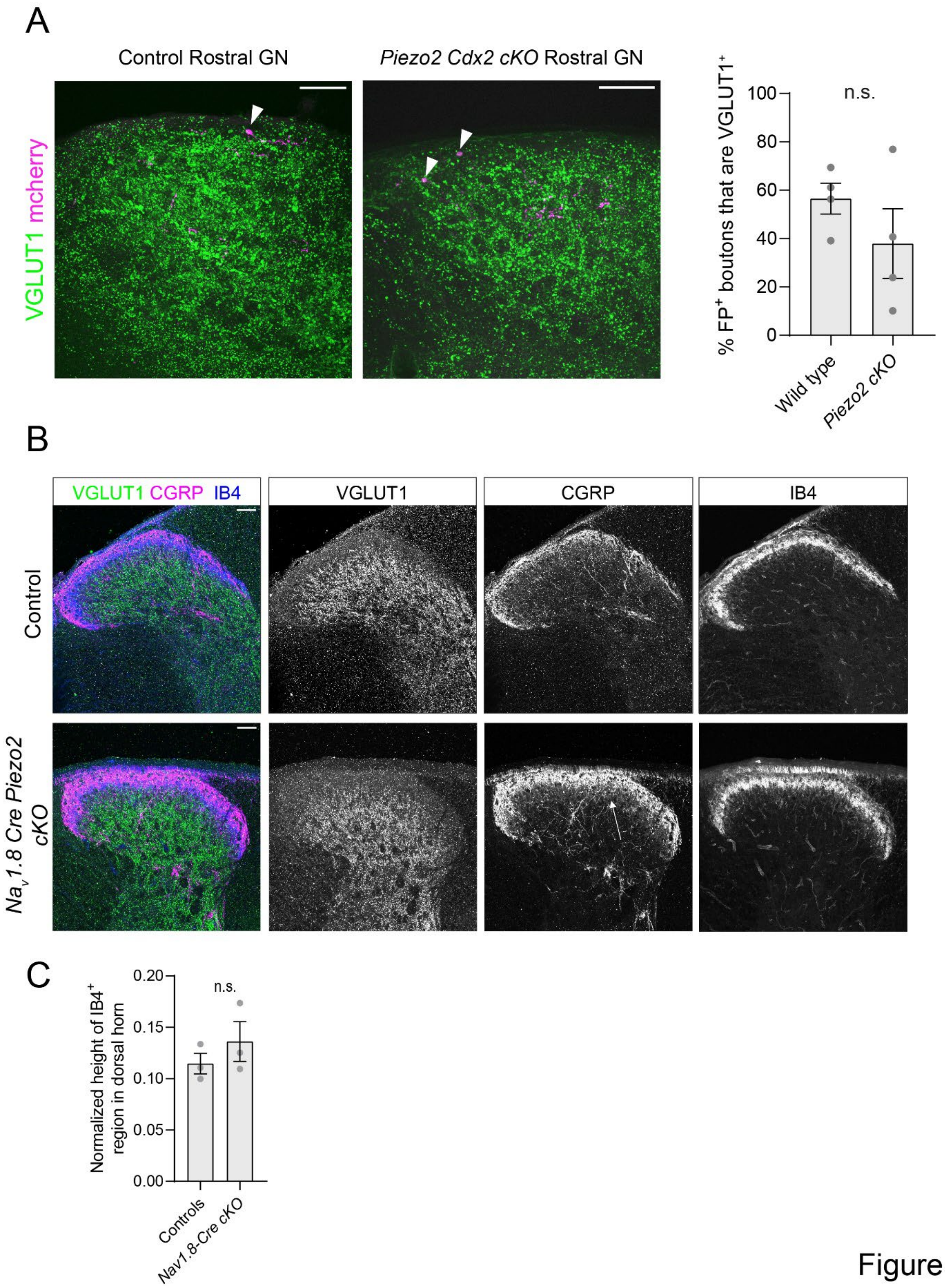
A
Control Rostral GN
Piezo2 Cdx2 cKO Rostral GN
VGLUT1 mcherry
% FP+ boutons that are VGLUT1+
n.s.
100
80
60
40
20
0
Wild type
Piezo2 cKO
B
VGLUT1 CGRP IB4
VGLUT1
CGRP
IB4
Control
Nav1.8 Cre Piezo2 cKO
C
Normalized height of IB4+ region in dorsal horn
n.s.
0.20
0.15
0.10
0.05
0.00
Controls
Nav1.8-Cre cKO

Figure S4

**Supplemental Figure 4: Additional characterization of somatosensory neuron central targeting in *Cdx2-Cre Piezo2 cKO* and *$Na_V1.8$-Cre Piezo2 cKO* mutants, related to Figure 4**

**A.** Left: Sections from gracile nuclei of control or *Cdx2-Cre/+; $Piezo2^{flox/Null}$* animals that were injected with AAV viruses to hind paw skin to sparsely label Aβ-LTMRs. Brainstem sections were stained with anti-VGLUT1 (Green) and anti-mcherry (magenta) antibodies to detect presynaptic inputs from all Aβ-LTMRs (VGLUT1) or from the sparsely labeled neurons (mcherry). In both genotypes, axons from Aβ-LTMRs ascended the dorsal column, innervated the gracile nuclei, and formed $VGLUT1^{+}$ presynaptic boutons (arrowheads). Scale bar = 50 µm. Right: Quantification of the percent of fluorescently labeled presynaptic boutons formed by hind paw Aβ-LTMRs in the brainstem that are positive for VGLUT1. There was no significant difference between control and *Piezo2 cKO* animals (unpaired t test). Each data point indicates one animal; averages are plotted; error bars indicate the s.e.m. "Wild type" refers to *+/+; $Piezo2^{+/+}$* animals and are a subset of the wild type animals that were used for quantification of Aβ-LTMR anatomy in [37]. "*Piezo2 cKO*" refers to *Cdx2-Cre/+; $Piezo2^{flox/Null}$* (n=2) and *Cdx2-Cre/+; $Piezo2^{flox/flox}$* (n=2) animals.

**B.** Example images of thoracic-level spinal cord sections from a control or *$Na_V1.8$-Cre; $Piezo2^{flox/Null}$* animal. One half of the dorsal horn of the spinal cord is shown. Spinal cords were stained using anti-VGLUT1 (green), anti-CGRP (purple), and for IB4-binding (blue). In *$Na_V1.8$-Cre; Piezo2 cKO* mutants, there is no detectable change in the height of the VGLUT1-negative region of the dorsal horn or in the height of the $IB4^{+}$ region. In contrast, the $CGRP^{+}$ region projects more deeply into the spinal cord (arrow). Scale bar = 50 µm.

**C.** Quantification of the height of the $IB4^{+}$ region in the dorsal horn of the spinal cord, normalized to the height of the dorsal horn. In *$Na_V1.8$-Cre; Piezo2 cKO* mutants, the $IB4^{+}$ region was not different than in control littermates (unpaired t-test). Each data point indicates one animal; averages are shown by horizontal bars; error bars indicate the s.e.m. Animals are the same as those used for the quantification shown in Figure 4E-F.

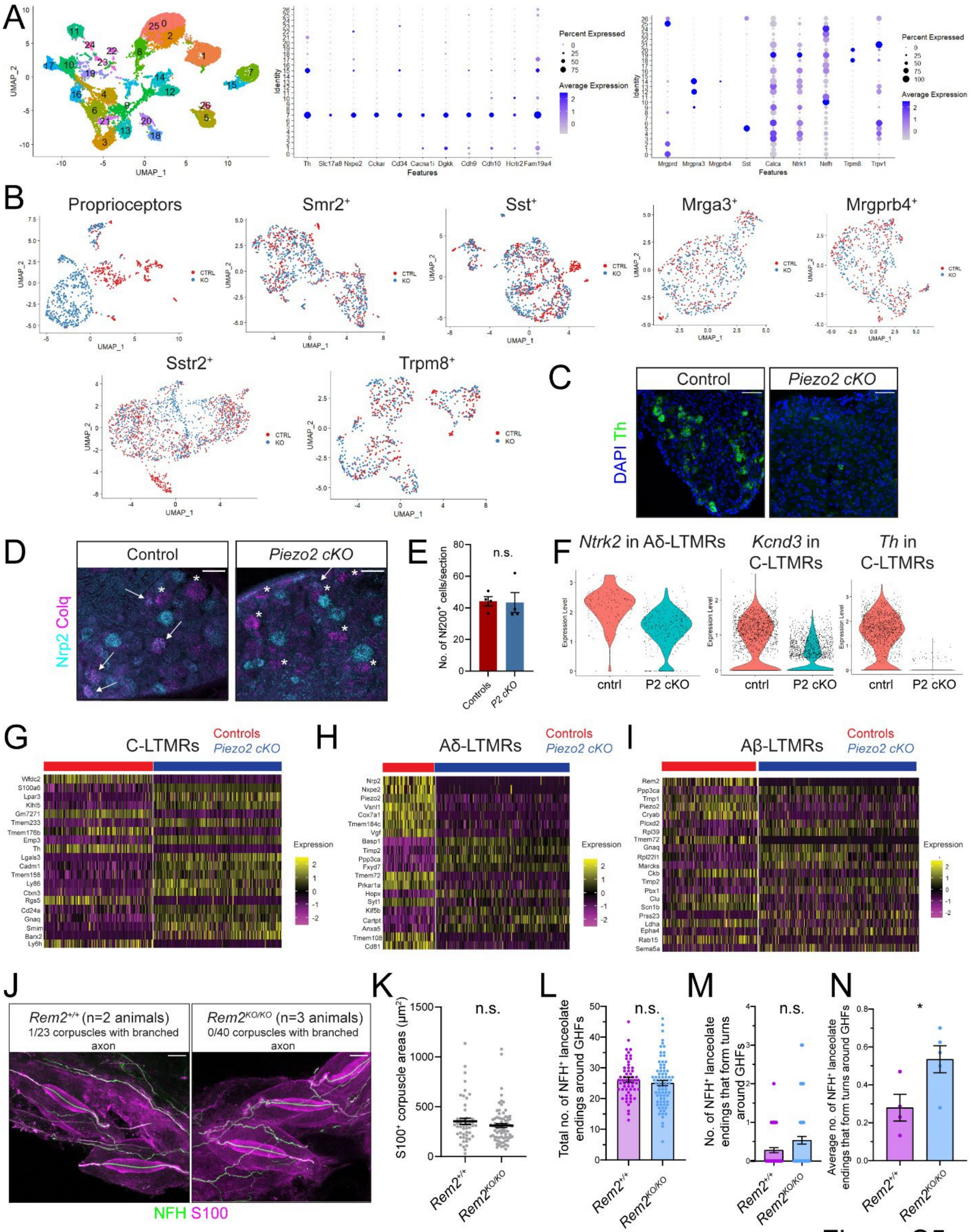
A
UMAP_1
UMAP_2
Identity
Features
Th Slc17a8 Nxpe2 Cckar Cd34 Cacna1i Dgkk Cdh9 Cdh10 Hcrtr2 Fam19a4
Percent Expressed
Average Expression
Mrgprd Mrgpra3 Mrgprb4 Sst Calca Ntrk1 Nefh Trpm8 Trpv1
B
Proprioceptors
Smr2⁺
Sst⁺
Mrga3⁺
Mrgprb4⁺
Sstr2⁺
Trpm8⁺
CTRL
KO
C
Control
Piezo2 cKO
DAPI Th
D
Control
Piezo2 cKO
Nrp2 Colq
E
n.s.
No. of Nf200⁺ cells/section
Controls
P2 cKO
F
Ntrk2 in Aδ-LTMRs
Kcnd3 in C-LTMRs
Th in C-LTMRs
Expression Level
cntrl
P2 cKO
G
C-LTMRs
Controls
Piezo2 cKO
Wfdc2
S100a6
Lpar3
Klhl5
Gm7271
Tmem233
Tmem176b
Emp3
Th
Lgals3
Cadm1
Tmem158
Ly86
Ctxn3
Rgs5
Cd24a
Gnaq
Smim
Barx2
Ly6h
Expression
H
Aδ-LTMRs
Controls
Piezo2 cKO
Nrp2
Nxpe2
Piezo2
Vsnl1
Cox7a1
Tmem184c
Vgf
Basp1
Timp2
Ppp3ca
Fxyd7
Tmem72
Prkar1a
Hopx
Syt1
Kif5b
Cartpt
Anxa5
Tmem108
Cd81
Expression
I
Aβ-LTMRs
Controls
Piezo2 cKO
Rem2
Ppp3ca
Timp1
Piezo2
Cryab
Plcxd2
Rpl39
Tmem72
Gnaq
Rpl22l1
Marcks
Ckb
Timp2
Pbx1
Clu
Scn1b
Prss23
Ldha
Epha4
Rab15
Sema5a
Expression
J
Rem2⁺/⁺ (n=2 animals)
1/23 corpuscles with branched axon
Rem2KO/KO (n=3 animals)
0/40 corpuscles with branched axon
NFH S100
K
n.s.
S100⁺ corpuscle areas (μm²)
Rem2⁺/⁺
Rem2KO/KO
L
n.s.
Total no. of NFH⁺ lanceolate endings around GHFs
Rem2⁺/⁺
Rem2KO/KO
M
n.s.
No. of NFH⁺ lanceolate endings that form turns around GHFs
Rem2⁺/⁺
Rem2KO/KO
N
*
Average no. of NFH⁺ lanceolate endings that form turns around GHFs
Rem2⁺/⁺
Rem2KO/KO

Figure S5

**Supplemental Figure 5: Additional characterization of single cell RNA sequencing data in *Cdx2-Cre Piezo2 cKO* animals, related to Figure 5**

**A.** Left: UMAP plot of DRG neurons collected from juvenile (P21-P24) control and *Cdx2-Cre; Piezo2 cKO* animals that met QC criteria after single cell RNA sequencing. Neurons are color-coded and labeled (0-26) by their computationally defined cluster identities. Middle: Dot plots showing expression levels of C-LTMR markers in each computationally defined cluster. Right: Dot plots of expression levels of genes that define other cell types.

**B.** UMAP plots of neurons corresponding to proprioceptors (clusters 11 and 24), *Smr2*$^{+}$/*Calca*$^{+}$/*Nefh*$^{+}$Aδ-HTMRs (cluster 4), SST$^{+}$ neurons (clusters 5 and 26), *Mrgpra3*$^{+}$/*Calca*$^{+}$ neurons (cluster 12), *Mrgpra3*$^{+}$/*Mrgprb4*$^{+}$ neurons (cluster 14), *Sstr2*$^{+}$/*Calca*$^{+}$ neurons (cluster 3), and *Trpm8*$^{+}$ neurons (clusters 18 and 20). Neurons corresponding to each cell type were subsetted from the main dataset and re-analyzed. Control (red) and *Piezo2 cKO* (blue) neurons are color-coded by genotype.

**C.** Representative images of lumbar-level DRG sections from control or *Cdx2-Cre; Piezo2 cKO* animals stained with RNAscope probes to detect *Th* (green) transcripts. DAPI staining is shown in blue. Scale bar = 50 µm. The stain was repeated in one additional control animal and one additional *Cdx2-Cre Piezo2 cKO* mutant, with similar results.

**D.** Representative images of thoracic-level DRG sections from control or *Cdx2-Cre; Piezo2 cKO* animals stained with RNAscope probes to detect *Colq* (purple, a marker for Aδ-LTMRs) or *Nrp2* (cyan) transcripts. Scale bar = 50 µm. Arrows point to *Colq*$^{+}$/*Nrp2*$^{+}$ cells, and asterisks indicate *Colq*$^{+}$/*Nrp2*$^{-}$ cells. The stain was repeated in one additional control animal and two additional *Cdx2-Cre Piezo2 cKO* mutants, with similar results.

**E.** Quantification of the average number of NF-200$^{+}$ cells in thoracic DRG sections for control or *Cdx2-Cre; Piezo2 cKO* animals. There was no difference between the two groups (Mann-Whitney U test). Each data point indicates one animal; averages are shown by horizontal bars; error bars indicate the s.e.m.

**F.** Violin plots showing expression levels of *Ntrk2* (encodes the TrkB receptor), *Kcnd3* (encodes $K_V$4.3), or Th transcripts in control and *Piezo2 cKO* Aδ-LTMRs or C-LTMRs, which were subsetted from the main RNA sequencing dataset.

**G-I.** Heat maps showing the top 20 most significantly differentially expressed genes when comparing control and *Cdx2-Cre; Piezo2 cKO* C-LTMRs (G), Aδ-LTMRs (H), or Aβ-LTMRs (I).

**J.** Images of Pacinian corpuscles from *Rem2*$^{+/+}$ or *Rem2*$^{-/-}$ animals. Tissue was stained using antibodies against NFH (green) and S100 (magenta). There was no increase in Pacinian axon branching in *Rem2* mutants. Scale bar = 50 µm. Data were collected from the following animals: *Rem2*$^{+/+}$ (n=2), *Rem2*$^{-/-}$ (n=3).

**K.** Quantification of the S100$^{+}$ areas of Meissner corpuscles in *Rem2*$^{+/+}$ or *Rem2*$^{-/-}$ animals. There was no significant difference between the two groups (Mann-Whitney U test). Each data point indicates one Meissner corpuscle; averages are shown by horizontal bars; error bars indicate the s.e.m. Data were collected from the following animals: *Rem2*$^{+/+}$ (n=3), *Rem2*$^{-/-}$ (n=3).

**L.** Quantification of NFH$^{+}$ axonal structures formed by lanceolate endings around guard hairs. There was no difference in the total number of NFH$^{+}$ lanceolate endings formed in *Rem2*$^{-/-}$ *KO* animals compared to controls (unpaired t test). Each data point indicates one hair follicle; averages are plotted; error bars indicate the s.e.m. Data were collected from the following animals: *Rem2*$^{+/+}$ (n=4), *Rem2* $^{-/-}$ (n=5).

**M.** Quantification of the total number of turns formed by NFH$^{+}$ lanceolate endings around guard hair follicles. There was no difference between *Rem2*$^{-/-}$*KO* animals compared to controls (Mann-Whitney U test). Each data point indicates one hair follicle; averages are plotted; error bars indicate the s.e.m. Data were collected from the same animals as in Supplemental Figure 5L.

**N.** Quantification of the average number of turns formed by NFH$^{+}$ lanceolate endings around guard hairs. There was a slight increase in *Rem2*$^{-/-}$*KO* animals compared to controls (*p<0.05, unpaired t test). Each data point indicates one animal; averages are plotted; error bars indicate the s.e.m. Data were collected from the same animals as in Supplemental Figure 5L-M.

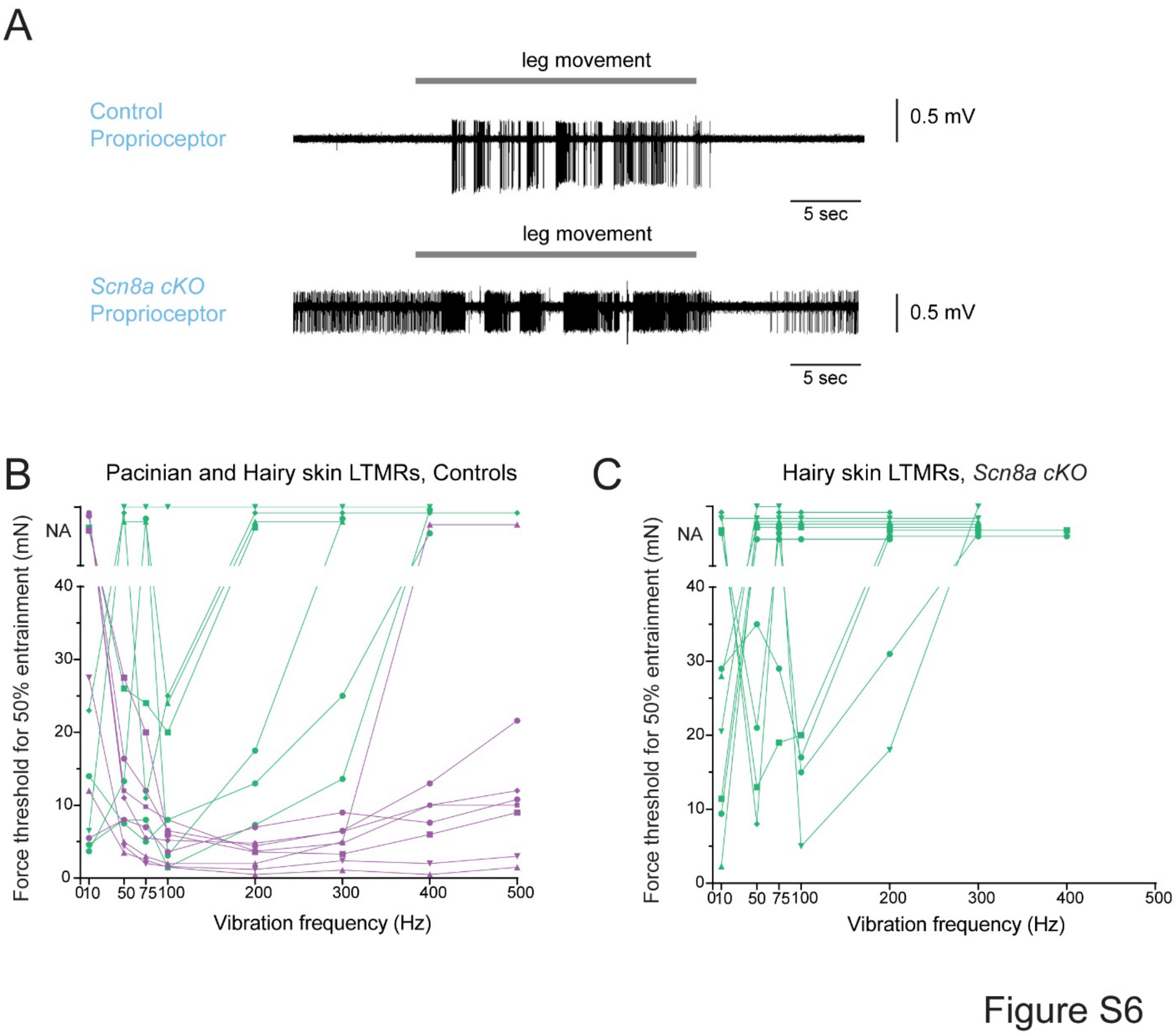

**Supplemental Figure 6: Additional characterization of Aβ-LTMR electrophysiological responses in *Avil*$^{Cre}$ *Scn8a*$^{flox}$ *cKO* animals and controls, related to Figure 6**

**A.** *In vivo* recording of proprioceptors from a control animal (top)(+/+*; Scn8a*$^{flox/+}$) or from an *Avil*$^{Cre/+}$*; Scn8a*$^{flox/flox}$ *cKO* animal (bottom). The gray rectangle indicates the time window during which the position of the hind limb was moved. **B.** The minimum force required for a unit to reach 50% entrainment (i.e., firing at least once every other cycle) when responding to a vibration stimulus is plotted for control animals. Pacinian units are in purple; hairy skin LTMRs are in green. "NA" indicates that a unit did not reach 50% entrainment at forces up to 35 mN (higher forces were not tested). Each data point indicates one unit tested at a specific frequency; points indicating the same neuron are connected by lines. **C.** The minimum force required for a hairy skin LTMR unit to reach 50% entrainment (i.e., firing at least once every other cycle) when responding to a vibration stimulus is plotted for *Scn8a cKO* animals. "NA" indicates that a unit did not reach 50% entrainment at forces up to 35 mN (higher forces were not tested). Each data point indicates one unit tested at a specific frequency; points indicating the same neuron are connected by lines.

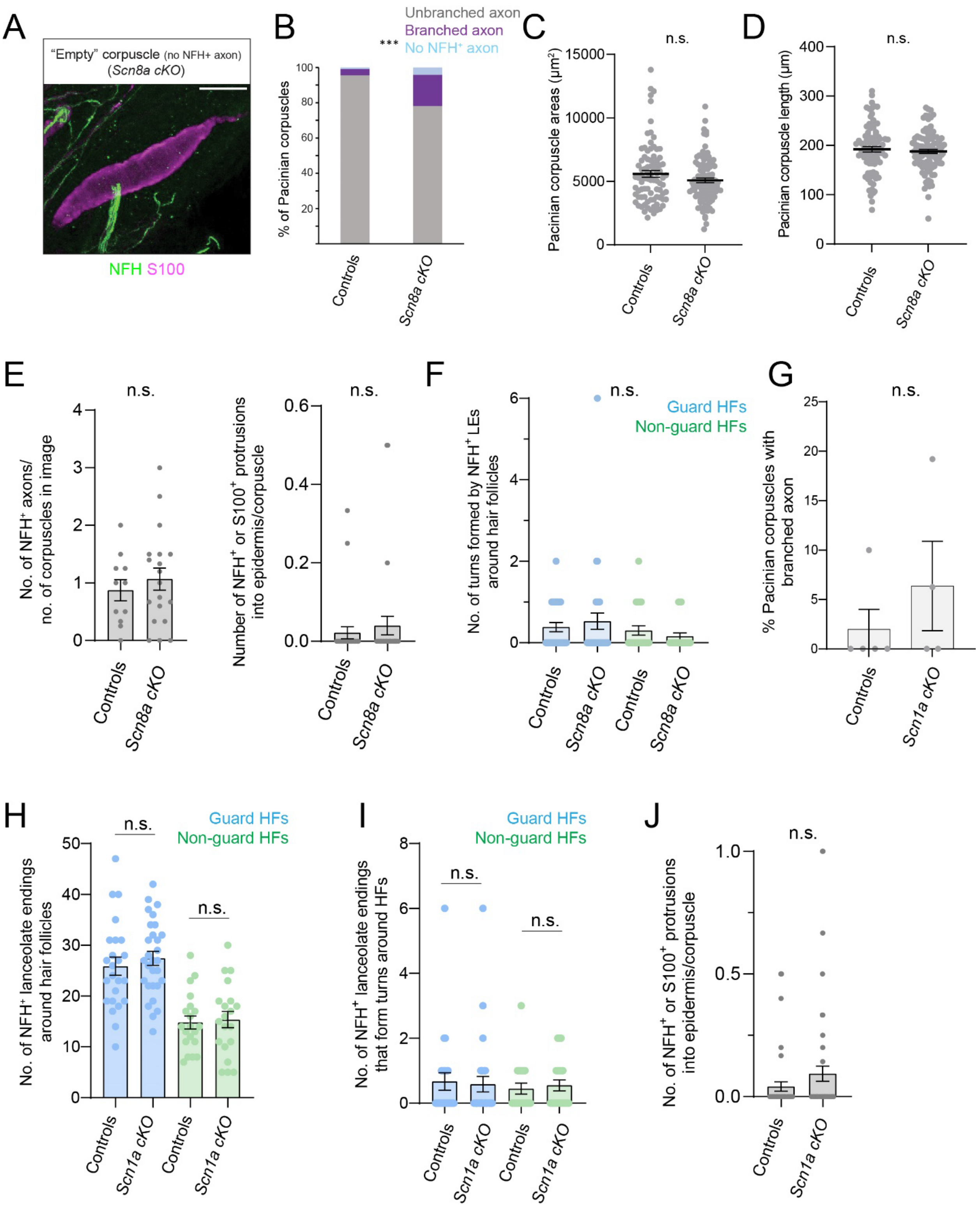

A
"Empty" corpuscle (no NFH+ axon)
(Scn8a cKO)
NFH S100
B
Unbranched axon
Branched axon
No NFH+ axon
% of Pacinian corpuscles
Controls
Scn8a cKO
C
n.s.
Pacinian corpuscle areas (µm²)
Controls
Scn8a cKO
D
n.s.
Pacinian corpuscle length (µm)
Controls
Scn8a cKO
E
n.s.
No. of NFH+ axons/ no. of corpuscles in image
Number of NFH+ or S100+ protrusions into epidermis/corpuscle
Controls
Scn8a cKO
F
n.s.
Guard HFs
Non-guard HFs
No. of turns formed by NFH+ LEs around hair follicles
Controls
Scn8a cKO
G
n.s.
% Pacinian corpuscles with branched axon
Controls
Scn1a cKO
H
n.s.
Guard HFs
Non-guard HFs
No. of NFH+ lanceolate endings around hair follicles
Controls
Scn1a cKO
I
n.s.
Guard HFs
Non-guard HFs
No. of NFH+ lanceolate endings that form turns around HFs
Controls
Scn1a cKO
J
n.s.
No. of NFH+ or S100+ protrusions into epidermis/corpuscle
Controls
Scn1a cKO

Figure S7

**Supplemental Figure 7: Additional characterization of Aβ-LTMR anatomies in *Avil*$^{Cre}$ *Scn8a*$^{flox}$ *cKO* animals and *Avil*$^{Cre}$ *Scn1a*$^{flox}$ *cKO* animals, related to Figure 7**

**A.** Example image of a Pacinian corpuscle from an *Avil*$^{Cre/+}$; *Scn8a*$^{flox/flox}$ animal that lacks an NFH$^{+}$ axon. Tissue was stained using antibodies against NFH (green) and S100 (magenta). Scale bar = 50 µm.

**B.** Pacinian corpuscles were categorized based on the presence and branching pattern of the NFH$^{+}$ axon inside the corpuscle. The proportions of corpuscles falling into the three categories were significantly different between control and *Avil*$^{Cre/+}$; *Scn8a*$^{flox/flox}$ animals (**$p<0.001$, Fisher's exact test).

**C.** Pacinian corpuscle areas in controls or *Avil*$^{Cre/+}$; *Scn8a*$^{flox/flox}$ animals were not significantly different (Mann-Whitney U test). Each data point indicates one corpuscle; averages are shown; error bars indicate the s.e.m. Data were collected from 8 controls and 8 *Avil*$^{Cre/+}$; *Scn8a*$^{flox/flox}$ mutants, which are a subset of the animals analyzed in Figure 7A.

**D.** Pacinian corpuscle lengths in controls or *Avil*$^{Cre/+}$; *Scn8a*$^{flox/flox}$ animals were not significantly different (Welch's t test). Each data point indicates one corpuscle; averages are shown; error bars indicate the s.e.m. Data were collected from the same animals as in Supplemental Figure 7C.

**E.** Left: Quantification of the number of NFH$^{+}$ axons innervating each Meissner corpuscle divided by the number of Meissner corpuscles present in an image. There was no significant difference between controls and *Avil*$^{Cre/+}$; *Scn8a*$^{flox/flox}$ animals (unpaired t test). Each data point indicates one tissue section image; averages are shown by horizontal bars; error bars indicate the s.e.m. Right: Quantification of the number of S100$^{+}$ or NFH$^{+}$ protrusions from Meissner corpuscles projecting into the epidermis, divided by the number of corpuscles detected in each tissue section. There was no significant difference between controls and *Avil*$^{Cre/+}$; *Scn8a*$^{flox/flox}$ animals (Mann-Whitney U test). Each data point indicates one image; averages are shown by horizontal bars; error bars indicate the standard error of the mean (s.e.m.). Data were collected from the same animals as in Figure 7C-D.

**F.** Quantification of NFH$^{+}$ axonal structures formed by lanceolate endings around guard hair follicles (blue) and non-guard hair follicles (green). There was no difference in the total number of turns formed by NF200$^{+}$ lanceolate endings between controls and *Avil*$^{Cre/+}$; *Scn8a*$^{flox/flox}$ animals, for either type of hair follicle (Mann-Whitney U test). Each data point indicates one hair follicle; averages are plotted; error bars indicate the s.e.m. Data were collected from the same animals as in Figure 7E.

**G.** Quantification of the percentage of Pacinian corpuscles with branched axons from adult animals. Each data point indicates one animal; averages are plotted; error bars indicate the s.e.m. There was no difference in Pacinian axon branching between control littermates and *Scn1a cKO* animals (Mann-Whitney U test).

**H.** Quantification of NFH$^{+}$ axonal structures formed by lanceolate endings around guard hair follicles (blue) and non-guard hair follicles (green). There was no difference in the total number of NFH$^{+}$ lanceolate endings between controls and *Avil*$^{Cre/+}$; *Scn1a*$^{flox/flox}$ animals, for either type of hair follicle (Mann-Whitney U test). Each data point indicates one hair follicle; averages are plotted; error bars indicate the s.e.m. Data were collected from 3 control animals and 3 *AvilCre/+; Scn1a$^{flox/flox}$* mutants.

**I.** Quantification of NFH$^{+}$ axonal structures formed by lanceolate endings around guard hair follicles (blue) and non-guard hair follicles (green). There was no difference in the number of turns formed by NFH$^{+}$ lanceolate endings between controls and *Avil$^{Cre/+}$; Scn1a$^{flox/flox}$* animals, for either type of hair follicle (unpaired t-test). Each data point indicates one hair follicle; averages are plotted; error bars indicate the s.e.m. Data were collected from the same animals as in Supplemental Figure 7H.

**J.** Quantification of the number of S100$^{+}$ or NFH$^{+}$ protrusions from Meissner corpuscles projecting into the epidermis, divided by the number of corpuscles detected in each tissue section. There was no significant difference between controls and *Avil$^{Cre/+}$; Scn1a$^{flox/flox}$* animals (Mann-Whitney U test). Each point indicates one image of a tissue section; averages are shown by horizontal bars; error bars indicate the standard error of the mean (s.e.m.). Data were collected from the same animals as Figure 7G-I.

**Supplemental Table 1.** List of all the genotypes and number of mice used for each experiment.

| **Experiment described in:** | **Control genotypes (number of animals)** | **Mutant genotypes (number of animals)** |
|---|---|---|
| Figure 1A | *+/+; Piezo2$^{flox/+}$* (n=3) | *Cdx2-Cre/+; Piezo2$^{flox/flox}$* (n=3) |
| | +/+; *Piezo2 $^{+/+}$* (n=1) | *Cdx2-Cre/+; Piezo2$^{flox/Null}$* (n=4) |
| | *Cdx2-Cre/+; Piezo2$^{+/+}$* (n=1) | |
| Figure 1B | *+/+; Piezo2$^{flox/flox}$* (n=1) | *Cdx2-Cre/+; Piezo2$^{flox/flox}$* (n=3) |
| | *+/+; Piezo2$^{flox/Null}$* (n=5) | *Cdx2-Cre/+; Piezo2$^{flox/Null}$* (n=2) |
| | *+/+; Piezo2$^{flox/+}$* (n=1) | |
| Figure 1E, Tuj1$^{+}$ quantification | +/+; *Piezo2$^{flox/+}$* (n=4) | *Cdx2-Cre/+; Piezo2$^{flox/flox}$* (n=4) |
| Figure 1E, NF200$^{+}$ quantification | *Piezo2$^{flox/flox}$* (n=3) | *Cdx2-Cre/+; Piezo2$^{flox/flox}$* (n=3) |
| | *Piezo2$^{flox/Null}$* (n=1) | *Cdx2-Cre/+; Piezo2$^{flox/Null}$* (n=1) |
| Figure 1F | *Piezo2$^{flox/flox}$* (n=3) | *Cdx2-Cre/+; Piezo2$^{flox/flox}$* (n=3) |
| | *Piezo2$^{flox/Null}$* (n=2) | *Cdx2-Cre/+; Piezo2$^{flox/Null}$* (n=2) |
| Figure 2B, P3-P5 | *+/+; Piezo2$^{flox/+}$* (n=2) | *Cdx2-Cre/+; Piezo2$^{flox/flox}$* (n=1) |
| | +/+; *Piezo2$^{flox/flox}$* (n=1) | *Cdx2-Cre/+; Piezo2$^{flox/Null}$* (n=2) |
| | +/+; *Piezo2$^{Null/+}$* (n=1) | |
| Figure 2B, P9-P10 | *+/+; Piezo2$^{flox/+}$* (n=2) | *Cdx2-Cre/+; Piezo2$^{flox/flox}$* (n=3) |
| | *+/+; Piezo2$^{Null/+}$* (n=1) | |
| Figure 2B, P14-P15 | *+/+; Piezo2$^{flox/+}$* (n=1) | *Cdx2-Cre/+; Piezo2$^{flox/flox}$* (n=2) |
| | *+/+; Piezo2$^{flox/flox}$* (n=1) | *Cdx2-Cre/+; Piezo2$^{flox/Null}$* (n=1) |
| | *+/+; Piezo2$^{flox/Null}$* (n=1) | |
| Figure 2D-F, P9 | *+/+; Piezo2$^{flox/+}$* (n=1) | *Cdx2-Cre/+; Piezo2$^{flox/Null}$* (n=3). |
| | *+/+; Piezo2$^{Null/+}$* (n=1) | |
| | *+/+; Piezo2$^{+/+}$* (n=1) | |
| Figure 2D-F, P14-P15 | *+/+; Piezo2$^{flox/flox}$* (n=4) | *Cdx2-Cre/+; Piezo2$^{flox/flox}$* (n=3) |
| | *+/+; Piezo2$^{flox/+}$* (n=1) | *Cdx2-Cre/+; Piezo2$^{flox/Null}$* (n=1) |
| | *Cdx2-Cre/+; Piezo2$^{Null/+}$* (n=1) | |
| Figure 3D-E, Supplemental Figure 3B | +/+; *Piezo2$^{flox/Null}$* (n=3) | *Dhh$^{Cre/+}$; Piezo2$^{flox/Null}$* (n=4) |

| | | |
|---|---|---|
| Figure 3F | *+/+; Piezo2*$^{flox/Null}$ (n=5) | *Dhh*$^{Cre/+}$*; Piezo2*$^{flox/Null}$ (n=8) |
| Figure 3G | *+/+; Piezo2*$^{flox/Null}$ (n=3) | *Dhh*$^{Cre/+}$*; Piezo2*$^{flox/Null}$ (n=3) |
| Figure 3H, Supplemental Figure 3G-H | *+/+; Piezo2*$^{flox/Null}$ (n=5) | *Dhh*$^{Cre/+}$*; Piezo2*$^{flox/Null}$ (n=5) |
| | | *Cdx2-Cre/+; Piezo2*$^{flox/flox}$ (n=3) |
| | | *Cdx2-Cre/+; Piezo2*$^{flox/Null}$ (n=2) |
| Figure 4B-C | *Cdx2-Cre/+; Piezo2*$^{+/+}$ (n=1) | *Cdx2-Cre/+; Piezo2*$^{flox/Null}$ (n=1) |
| | *Piezo2*$^{flox/flox}$ (n=4) | *Cdx2-Cre/+; Piezo2*$^{flox/flox}$ (n=4) |
| Figure 4D | *Cdx2-Cre/+; Piezo2*$^{+/+}$ (n=1) | *Cdx2-Cre/+; Piezo2*$^{flox/Null}$ (n=1) |
| | *+/+; Piezo2*$^{flox/flox}$ (n=4) | *Cdx2-Cre/+; Piezo2*$^{flox/flox}$ (n=3) |
| Figure 4E-F, Supplemental Figure 4B | *+/+; Piezo2*$^{flox/Null}$ (n=3) | *Na*$_{v}$*1.8-Cre/+; Piezo2*$^{flox/Null}$ (n=3) |
| Figure 5A-G (RNAseq data) | *+/+; Piezo2*$^{flox/+}$ (n=1) | *Cdx2-Cre/+; Piezo2*$^{flox/flox}$ (n=5) |
| | *+/+; Piezo2*$^{flox/flox}$ (n=2) | |
| | *+/+; Piezo2*$^{Null/+}$ (n=1) | |
| | *Cdx2-Cre/+; Piezo2*$^{flox/+}$ (n=1) | |
| Figure 5J | *Cdx2-Cre/+; Piezo2*$^{+/+}$ (n=1) | *Cdx2-Cre/+; Piezo2*$^{flox/flox}$ (n=2) |
| | *+/+; Piezo2*$^{flox/flox}$ (n=2) | *Cdx2-Cre/+; Piezo2*$^{flox/Null}$ (n=1). |
| Figure 5K | *Cdx2-Cre/+; Piezo2*$^{+/+}$ (n=1) | *Cdx2-Cre/+; Piezo2*$^{flox/flox}$ (n=2) |
| | *+/+; Piezo2*$^{flox/flox}$ (n=2) | *Cdx2-Cre/+; Piezo2*$^{flox/Null}$ (n=1). |
| Figure 5L | *+/+; Piezo2*$^{flox/flox}$ (n=2) | *Cdx2-Cre/+; Piezo2*$^{flox/flox}$ (n=3) |
| | *Piezo2*$^{flox/+}$ (n=1) | |
| Figure 5M | *Cdx2-Cre/+; Piezo2*$^{+/+}$ (n=1) | *Cdx2-Cre/+; Piezo2*$^{flox/flox}$ (n=2) |
| | *+/+; Piezo2*$^{flox/flox}$ (n=2) | *Cdx2-Cre/+; Piezo2*$^{flox/Null}$ (n=2) |
| | *+/+; Piezo2*$^{flox/+}$ (n=1) | |
| Figure 5N | *Piezo2*$^{flox/flox}$ (n=2) | *Cdx2-Cre/+; Piezo2*$^{flox/flox}$ (n=3) |
| | *+/+; Piezo2*$^{flox/+}$ (n=2) | *Cdx2-Cre/+; Piezo2*$^{flox/Null}$ (n=1) |

| | | |
|---|---|---|
| Figure 5O | *Piezo2*$^{flox/flox}$ (n=1) | *Cdx2-Cre/+; Piezo2*$^{flox/flox}$ (n=3) |
| | +/+; *Piezo2*$^{flox/+}$ (n=2) | |
| Figure 6B | +/+; +/+ (n=3) | *Avil*$^{Cre/+}$; *Scn8a*$^{flox/flox}$ (n=7) |
| | +/+; *Scn8a*$^{flox/+}$ (n=5) | |
| | +/+; *Scn8a*$^{flox/flox}$ (n=2) | |
| Figure 7A | +/+; *Scn8a*$^{flox/+}$ (n=3) | *AvilCre/+; Scn8a*$^{flox/flox}$ (n=10) |
| | +/+; *Scn8a*$^{flox/flox}$ (n=5) | |
| Figure 7C-D, Supplemental Figure 7E | +/+; *Scn8a*$^{flox/+}$ (n=2) | *AvilCre/+; Scn8a*$^{flox/flox}$ (n=5) |
| | +/+; *Scn8a*$^{flox/flox}$ (n=3) | |
| Figure 7E, Supplemental Figure 7F | +/+; *Scn8a*$^{flox/flox}$ (n=3) | *AvilCre/+; Scn8a*$^{flox/flox}$ (n=4) |
| | +/+; *Scn1a*$^{flox/+}$; *Scn8a*$^{flox/+}$ (n=1) | |
| Figure 7G-I, Supplemental Figure 7J | +/+; *Scn1a*$^{flox/flox}$ (n=1) | *Avil*$^{Cre/+}$; *Scn1a*$^{flox/flox}$ (n=6) |
| | +/+; *Scn1a*$^{flox/+}$ (n=4) | |
| Supplemental Figure 1E | +/+; *Piezo2*$^{flox/flox}$ (n=1) | *Cdx2-Cre/+; Piezo2*$^{flox/flox}$ (n=2) |
| | +/+; *Piezo2*$^{flox/+}$ (n=1) | *Cdx2-Cre/+; Piezo2*$^{flox/Null}$ (n=1) |
| | +/+; *Piezo2*$^{flox/Null}$ (n=1) | |
| Supplemental Figure 1G | +/+; *Piezo2*$^{flox/+}$ (n=5) | *Cdx2-Cre/+; Piezo2*$^{flox/flox}$ (n=4) |
| | +/+; *Piezo2*$^{flox/Null}$ (n=1) | *Cdx2-Cre/+; Piezo2*$^{flox/Null}$ (n=1) |
| Supplemental Figure 1H | +/+; *Piezo2*$^{flox/+}$ (n=3) | *Cdx2-Cre/+; Piezo2*$^{flox/flox}$ (n=3) |
| Supplemental Figure 2B-C, P5 | +/+; *Piezo2*$^{flox/flox}$ (n=1) | *Cdx2-Cre/+; Piezo2*$^{flox/flox}$ (n=1) |
| | +/+; *Piezo2*$^{flox/+}$ (n=1) | *Cdx2-Cre/+; Piezo2*$^{flox/Null}$ (n=1) |
| Supplemental Figure 2B-C, P9 | +/+; *Piezo2*$^{flox/+}$ (n=2) | *Cdx2-Cre/+; Piezo2*$^{flox/flox}$ (n=3) |
| Supplemental Figure 2B-C, P15 | +/+; *Piezo2*$^{flox/flox}$ (n=1) | *Cdx2-Cre/+; Piezo2*$^{flox/flox}$ (n=2) |
| | +/+; *Piezo2*$^{flox/+}$ (n=1) | |
| | +/+; *Piezo2*$^{flox/Null}$ (n=1) | |
| Supplemental Figure 2B-C, Adult | +/+; *Piezo2*$^{flox/flox}$ (n=2) | *Cdx2-Cre/+; Piezo2*$^{flox/flox}$ (n=6) |

| | | |
|---|---|---|
| | +/+; *Piezo2*$^{flox/Null}$ (n=4) | *Cdx2-Cre/+; Piezo2*$^{flox/Null}$ (n=2) |
| | +/+; *Piezo2*$^{flox/+}$ (n=2) | |
| Supplemental Figure 3F | +/+; *Piezo2*$^{flox/Null}$ (n=4) | *Dhh*$^{Cre/+}$*; Piezo2*$^{flox/Null}$ (n=5) |
| Supplemental Figure 4A | +/+; *Piezo2* $^{+/+}$ (n=4) | *Cdx2-Cre/+; Piezo2*$^{flox/flox}$ (n=2) |
| | | *Cdx2-Cre/+; Piezo2*$^{flox/Null}$ (n=2) |
| Supplemental Figure 5E | +/+; *Piezo2*$^{flox/flox}$ (n=3) | *Cdx2-Cre/+; Piezo2*$^{flox/flox}$ (n=4) |
| | +/+; *Piezo2*$^{flox/+}$ (n=1) | |
| Supplemental Figure 7C-D | +/+; *Scn8a*$^{flox/+}$ (n=4) | *AvilCre/+; Scn8a*$^{flox/flox}$ (n=8) |
| | +/+; *Scn8a*$^{flox/flox}$ (n=4) | |
| Supplemental Figure 7G | +/+; *Scn1a*$^{flox/+}$ (n=4) | *AvilCre/+; Scn1a*$^{flox/flox}$ (n=4) |
| | +/+; *Scn1a*$^{flox/flox}$ (n=1) | |
| Supplemental Figure 7H-I | +/+; *Scn1a*$^{flox/flox}$ (n=2) | *AvilCre/+; Scn1a*$^{flox/flox}$ (n=3) |
| | +/+; *Scn1a*$^{flox/+}$ (n=1) | |
